# Quantifying spatiotemporal decoupling of GDGT-temperature relationships in a deep alpine lake

**DOI:** 10.1101/2025.08.01.668065

**Authors:** Jingjing Li, Fanfan Meng, Qian Wang, B. David A. Naafs, Francien Peterse, Rong Wang, Huan Yang, Xiangdong Yang, Jianjun Wang, Shucheng Xie

**Author notes:** Corresponding author. E-mail address (Jianjun Wang).

## Abstract

Glycerol dialkyl glycerol tetraethers (GDGTs), membrane lipids produced by archaea and some bacteria, are widely used in paleoclimate reconstructions due to their empirical relationship with temperature. However, their application in lakes is complicated by uncertainties in source attribution and environmental controls on their distribution. To address these constraints, we analyzed both branched and isoprenoid GDGTs (brGDGTs and isoGDGTs) in settling particles collected over 19 months using sediment traps at four depths (10, 15, 25 and 35 m) in Lake Lugu, a deep, stratified alpine lake in southwestern China. GDGT fluxes showed synchronous spatiotemporal variations across depths, with higher values during winter mixing than summer stratification, suggesting in situ production enhanced by nutrient upwelling during lake overturn. Correlations between GDGT distributions and high-resolution water temperature profiles revealed strong temperature sensitivity in isoGDGTs, particularly the Ring Index (RI), with peak correlations linked to mean temperatures ∼20 days prior to trap recovery, indicating a clear temporal lag in GDGT–temperature relationship. Moreover, stronger correlations with temperatures at overlying depths, implying vertical transport of isoGDGTs and a dominant autochthonous origin from the upper water column. In contrast, brGDGTs displayed weak or non-significant temperature dependence, likely reflecting distinct microbial sources or other controlling factors. These findings underscore the utility of isoGDGT-based proxies, particularly RI, while highlighting the importance of accounting for spatiotemporal offsets in GDGTs production when reconstructing paleotemperatures in deep, stratified lakes.

## 1. Introduction

Glycerol dialkyl glycerol tetraethers (GDGTs) are membrane-spanning lipids synthesized by many archaea and certain bacteria (<u>Schouten et al., 2013)</u>. Isoprenoid GDGTs (isoGDGTs) are exclusively produced by archaea. These compounds comprise isoprenoid chains with a varying number of cyclopentane moieties, denoted as GDGT-0 to –8 according to the amount of cyclopentane moieties. Among them, the acyclic GDGT-0 is the most common isoGDGT and is synthesized by a broad range of archaea groups, including ammonia-oxidizing Thaumarchaeota, anaerobic methanotrophs, methanogens, and uncultured crenarchaeota (<u>Pancost et al., 2001</u>; <u>Blaga et al., 2009</u>; <u>Sinninghe Damsté et al., 2012a</u>; <u>Buckles et al., 2013</u>; <u>Li et al., 2019</u>). GDGT-1 to –3 are commonly found in Euryarchaeota, Crenarchaeota and Thaumarchaeota (<u>Schouten et al., 2013</u>). Notably, crenarchaeol, a unique isoGDGT containing four cyclopentane moieties and one cyclohexane moiety, is exclusively produced by Thaumarchaeota, along with its isomer (crenarchaeol’) (<u>SinningheDamsté et al., 2002</u>; <u>Elling et al., 2017</u>). The degree of cyclization of isoGDGTs in marine sediments correlates strongly with sea surface temperature, and this relationship underpins the widely used TEX_86_ (TetraEther indeX of 86 carbon atoms) paleotemperature proxy (<u>Schouten et al., 2002</u>). TEX_86_ has been broadly applied to reconstruct past sea surface temperatures for marine sedimentary records (<u>Taylor et al., 2013</u>; <u>Zhang et al., 2014</u>; <u>Tierney and Tingley, 2015</u>). Additionally, the Ring Index (RI), representing the weighted average number of cyclopentane moieties in isoGDGTs, has shown robust temperature sensitivity in both laboratory cultures and hot spring environments (<u>De Rosa et al., 1980</u>; <u>Kaur et al., 2015</u>).

Branched GDGTs (brGDGTs), a second group of GDGTs synthesized by bacteria, contain 4–6 methyl branches and up to two cyclopentane moieties on two alkyl chains (<u>Sinninghe Damsté et al., 2000</u>; <u>Weijers et al., 2006</u>). BrGDGTs are globally distributed and are particularly abundant in mineral soils, lakes, and peats (<u>Hopmanset al., 2004</u>; <u>Weijers et al., 2007</u>; <u>Naafs et al., 2017b</u>; <u>Martínez-Sosa et al., 2021</u>). Although the full range of their biological sources remain unidentified, *Acidobacteria* have been considered as key producers of brGDGTs (<u>Halamka et al., 2021</u>; <u>Chen et al., 2022</u>). The methylation and cyclization of brGDGTs (MBT/CBT) have been empirically associated with temperature and pH in globally distributed soils (<u>Weijerset al., 2007</u>; <u>Peterse et al., 2012</u>). Furthermore, the separation of 5-methyl and 6-methyl brGDGT isomers has revealed that only the 5-methyl brGDGTs are sensitive to temperature, while 6-methyl brGDGTs are more strongly associated with pH (<u>De Jonge et al., 2013</u>; <u>2014a</u>; <u>Yang et al., 2015</u>; <u>Naafs et al., 2017a</u>; <u>2017b</u>; <u>Dearing Crampton-Flood et al., 2020</u>).

The empirical relationships between GDGT distributions and environmental variables such as temperature and pH form the basis for their use in paleoclimate reconstructions across a range of sedimentary archives (<u>Tierney et al., 2008</u>; <u>Knies et al., 2014</u>; <u>Zhang et al., 2014</u>; <u>Johnson et al., 2016</u>; <u>Naafs et al., 2018</u>). However, applying GDGT-based proxies, particularly TEX_86_, in lacustrine environments poses challenges due to uncertainties in their sources and ecological controls. While isoGDGTs in lakes are generally associated with Thaumarchaeota in the water column, contributions from methanogens, methanotrophs, or soil-derived archaea may complicate the interpretation of temperature signal (<u>Blaga et al., 2009</u>; <u>Sinninghe Damsté et al., 2012a</u>; <u>Buckles et al., 2013</u>; <u>Li et al., 2019</u>; <u>Zheng et al., 2022</u>).

Moreover, the ecological niche of Thaumarchaeota, which typically reside in deeper, colder oxygenated waters, may result in TEX_86_ reflecting subsurface rather than surface temperatures (<u>Schouten et al., 2012</u>; <u>Zhang et al., 2016b</u>; <u>Kumar et al., 2019</u>; <u>Yao et al., 2019</u>; <u>Sinninghe Damsté et al., 2022</u>). In Lake Chala, Thaumarchaeotal blooms were suppressed under shallow oxycline conditions, leading to TEX_86_ signal that was biased due to limited production (<u>Baxter et al., 2021</u>). Similarly, elevated TEX_86_-derived temperatures observed under low dissolved oxygen (DO) conditions in a deep stratified lake further support this interpretation (<u>Zheng et al., 2022</u>).

The application of brGDGT-based proxies is also complicated by multiple factors. In situ production of brGDGTs within water columns and sediments challenges the validity of soil-based transfer functions (<u>Sinninghe Damsté et al., 2009</u>; <u>Tierney and Russell, 2009</u>; <u>Li et al., 2016</u>; <u>Colcord et al., 2017</u>; <u>Weber et al., 2018</u>; <u>van Bree et al., 2020</u>). As a result, lake-specific calibrations have been proposed (<u>Tierney et al., 2010</u>; <u>Pearson et al., 2011</u>; <u>Dang et al., 2018</u>; <u>Russell et al., 2018</u>; <u>Martínez-Sosa et al., 2021</u>; <u>Wu et al., 2023</u>; <u>Zhao et al., 2023</u>), which differ significantly from those developed for soils and peats (<u>Naafs et al., 2017a</u>; <u>2017b</u>).

These discrepancies suggest the presence of different source organisms or distinct environmental controls acting on aquatic brGDGT producers. Additionally, the relative contributions of aquatic-versus soil-derived brGDGTs can vary both between and within lakes on seasonal timescales, further complicating their use as temperature proxies. The biological sources and ecophysiology of brGDGT producers remain poorly constrained and likely differ across soils, lake sediments, and the water column (<u>Buckles et al., 2014</u>; <u>Weber et al., 2018</u>), adding to the uncertainty.

To better constrain the production and transfer dynamic of GDGTs in lakes, we investigated the temporal and vertical distribution of br– and isoGDGTs in settling particles from Lake Lugu, a deep, stratified alpine lake in southwest China (Fig. 1) (<u>Wang and Dou, 1998</u>). Sediment traps were deployed at four depths (10, 15, 25, and 35 m), and samples were collected at two-to three-month intervals from December 2012 to April 2014. This sampling strategy allowed for the assessment of seasonal and depth-dependent variability in GDGT fluxes. Concurrently, high-resolution water temperature profiles were recorded every 2 hours at 0.5 m intervals from 0 to 20 m, and every 2 m from 20 to 40 m during the 60 days preceding each trap recovery. By integrating GDGT distributions with spatiotemporal temperature data, we examined the temperature sensitivity of individual GDGTs and GDGT-based proxies, their vertical transport dynamics, and the temporal lag between GDGT production and export. These findings offer valuable insights into the interpretation of GDGT signals in lacustrine sedimentary archives and contribute to refining their use for paleoclimate reconstructions in stratified lake systems.

**Fig. 1.**
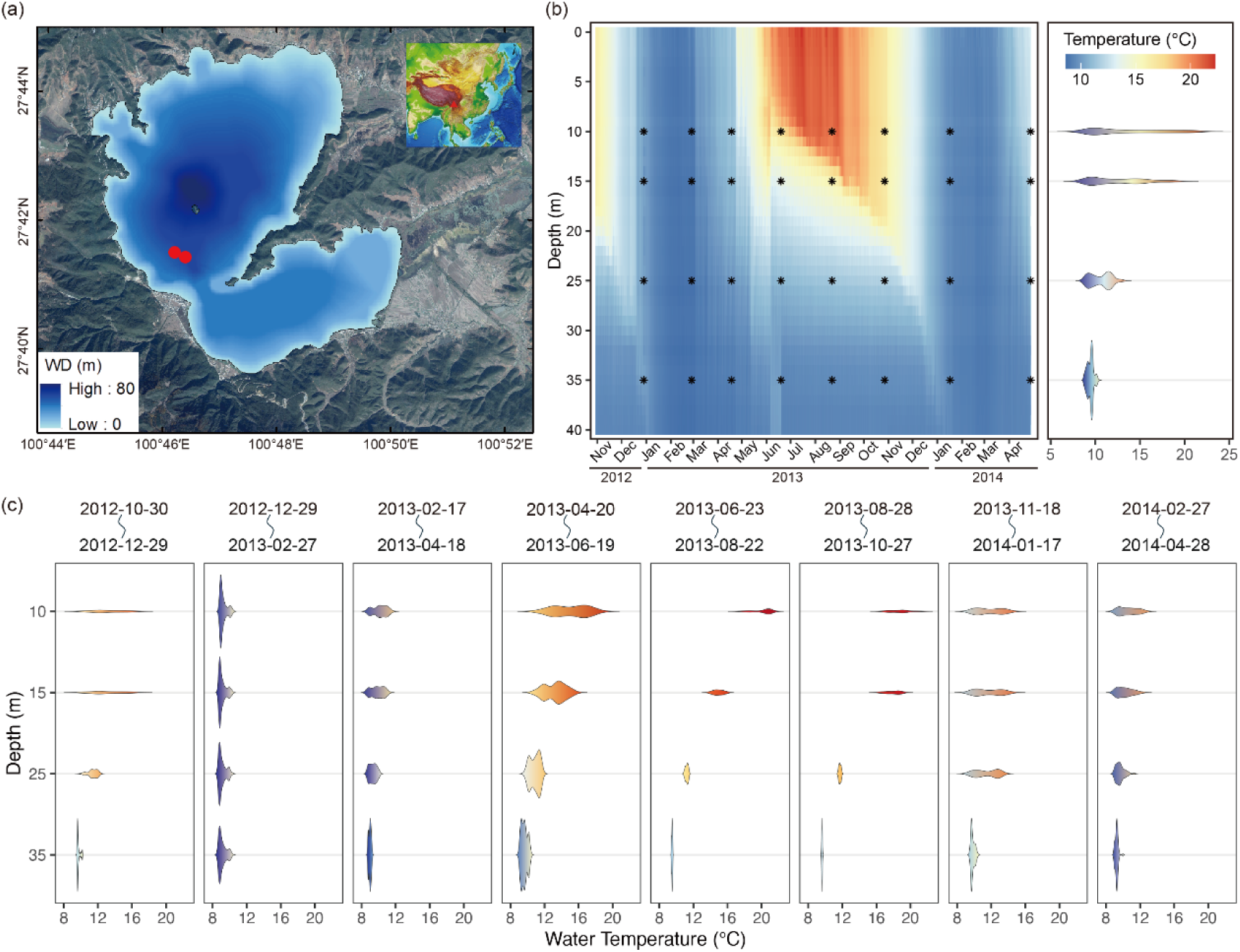
Sampling information and water column temperature variation in Lake Lugu from October 2012 to April 2014. (a) Sampling locations and trap deployment depths at two sites: TP1 (27°41′30.13″N, 100°46′17.23″E) and TP2 (27°41′24.74”N, 100°46′30.39″E), with traps suspended at 10 and 35 m (TP1); and 15 and 25 m (TP2). (b) High-resolution water temperature profiles at the four trap depths, with black asterisks indicating the deployment depths, along with the corresponding trap recovery dates. Sediment traps were initially deployed in October 2012, and subsequently emptied and redeployed on the same day at two-month intervals from December 2012 to October 2013, and at three-month intervals from January 2014 to April 2014. The yellow and blue shaded areas indicate periods of thermal stratification and mixing in the water column, respectively. Violin plots illustrating water temperature variations at each trap deployment depth. Plot (b) shows temperature variation over the entire monitoring period, while plot (c) indicates temperature variation during each of the eight sampling intervals (calculated over the 60 days prior to each trap recovery date).

## 2. Materials and methods

### 2.1. Study site

Lake Lugu (27°41′–27°45′N, 100°45′–100°50′E; 2691 m a.s.l.) is one of the deepest alpine freshwater lakes located on the southeastern margin of the Tibetan Plateau. It sits at the border of northwestern Yunnan and southwestern Sichuan provinces in southwest China (<u>Wang and Dou, 1998</u>) (Fig. 1a). The lake covers a surface area of 48.45 km^2^ and has a catchment of approximately 171.4 km^2^. Its mean and maximum depths are 40.3 and 93.5 m, respectively. Lake Lugu is a hydrologically semi-closed system with a water residence time of approximately 18 years. It is primarily sustained by precipitation, ephemeral runoff, and inflow from several small tributaries, including the Shankua and Sanjiacun Rivers, which exert limited hydrological influence (<u>NIGLAS, 2019</u>; <u>Chang et al., 2022</u>). The lake drains through a single outlet, the Gaizu River, which flows into the Yalong River via the Caohai wetland (<u>Yang, 1984</u>).

This lake consists of two sub-basins aligned northwest–southeast, separated by a prominent peninsula that extends across much of its center portion (<u>Zhang et al., 1997</u>) (Fig. 1a). It is a subtropic monomictic lake, with thermal stratification initiating in early April and intensifying from May through September. Stratification breaks down in early winter, and complete water column mixing typically occurs from January to March (<u>Wang et al., 2018</u>). During summer, seasonal stratification generates pronounced vertical gradients in DO, electrical conductivity, and pH, whereas winter mixing leads to a more homogenous vertical profile (<u>Chang et al., 2022</u>). The lake remains ice-free throughout the year and maintains high water transparency, with a maximum Secchi depth of 12–14 m. According to national environmental standards, the water quality of Lake Lugu is classified as Class I (<u>NIGLAS, 2019</u>). It currently exhibits oligotrophic conditions, with total phosphorus (TP) and total nitrogen (TN) concentrations of approximately 12.7 and 107.9 μg·l^−1^, respectively (<u>Wang et al., 2018</u>).

The regional climate is dominated by the Indian summer monsoon from June to September, and the southern branch of the mid-latitude westerlies in winter (<u>An et al., 2011</u>). Annual precipitation averages approximately 940 mm, with around 90% of it falling during the monsoon season (<u>Wang et al., 2014</u>). The mean annual air temperature is approximately 12.8 °C, with monthly averages ranging from 5.6 °C in December to 19.4 °C in July (<u>Wang et al., 2016</u>).

### 2.2. Sample collection

Sediment traps were deployed at two sites in the northern basin of Lake Lugu: TP1 (27°41′30.13″N, 100°46′17.23″E) and TP2 (27°41′24.74”N, 100°46′30.39″E), where the water depth is approximately 40 m (Fig. 1b). At each site, traps were suspended at two depths: 10 and 35 m at TP1, and 15 and 25 m at TP2 (<u>Wang et al., 2018</u>). The upper edge of the trap was positioned approximately 5 m above the lake bottom, as described in <u>Wang et al. (2018)</u>. Each trap consisted of four parallel transparent PVC tubes (50 cm length, 64 cm^2^ active area). Materials collected from all four tubes at each depth were pooled and homogenized for analysis. The sediment traps used in this study were part of the same sampling campaign as those in <u>Wang et al. (2018)</u>, but were used for different analytical purposes. Specifically, <u>Wang et al.</u> (<u>2018)</u> focused on Cladocera and diatom assemblages, while this study investigates lipid biomarkers from the remaining unpoisoned trap material.

Trap deployment began in October 2012, with subsequent retrieval and redeployment on the same day at bimonthly intervals from December 2012 to October 2013, and at trimonthly intervals from January 2014 to April 2014 (Fig. 1b and Table S2). The minimum deployment duration was approximately 60 days. Bulk mass fluxes (g·m^−2^·d^−1^) were calculated based on dry weight, deployment duration, and trap area. Although some in situ production of GDGTs may have occurred within the traps, it is generally assumed to be minimal, and is not explicitly quantified in this study.

Water column temperature was recorded every 2 hours from October 2012 to April 2014 using HOBO TidbiT v2 data loggers (Onset, USA), which were mounted on a vertically moored cable. Temperature sensors were spaced at 0.5 m intervals from 0 and 20 m, and at 2 m intervals from 20 to 40 m. Basic physicochemical parameters, including DO, pH, and conductivity were measured on each trap recovery date using a YSI 600XL multi-parameter sonde (Yellow Springs, USA), as detailed by <u>Wang et al. (2018)</u>. These data are not repeated here.

### 2.3. Lipid extraction

Collected materials were freeze-dried, homogenized, and weighted to determine particle fluxes (g·m^−2^·d^−1^). Lipids were extracted following the protocol described by <u>Li et al. (2022)</u>. Briefly, ∼0.2 g of dry sample was ultrasonically extracted four times (10 min each) with dichloromethane/methanol (9:1, v/v) mixture. The combined extract was saponified with 6% KOH in methanol at room temperature for 12 h. Neutral lipids were extracted with hexane, and separated via column chromatography over activated silica gel, using hexane for apolar fractions and methanol for polar fractions. The polar fraction containing core GDGTs was redissolved in hexane/isopropanol (99:1, v/v), filtered through a 0.45 μm PTFE filter, and dried under gentle nitrogen flow for instrumental analysis.

### 2.4. GDGT analysis

GDGTs were analyzed using high-performance liquid chromatography– atmospheric pressure chemical ionization–mass spectrometry (HPLC–APCI–MS) with triple quadruple detection on an Agilent 1200 series (USA), following the method of <u>Yang et al. (2015)</u>. Chromatographic separation was achieved using two Thermo Finnigan silica columns (150 mm × 2.1 mm, 1.9 μm) in tandem. The mobile phase consisted of a gradient from 84% A (*n*-hexane) and 16% B (ethyl acetate), held for 5 min, followed by a linear gradient to 82% A and 18% B over 60 min, then to 100% B over 21 min, held for 4 min, and returned to 84% A and 16% B to equilibrate the column for 30 min. The flow rate was 0.2 ml·min^-1^. Detection was performed in single ion monitoring (SIM) mode, targeting [M+H]^+^ ions: *m/z* 1302, 1300, 1298, 1296, and 1292 for isoGDGTs, and *m/z* 1050, 1048, 1046, 1036, 1034, 1032, 1022, 1020, and 1018 for brGDGTs. Quantification was based on the integration of peak areas relative to a synthetic internal standard (C_46_ GTGT, *m/z* 744, 10 μl of 11.57 ng·μl^-1^) (<u>Huguet et al., 2006</u>). Concentrations are considered semi-quantitative, assuming equal ionization efficiency and a 1:1 response factor between GDGTs and the internal standard. Recovery rates were not determined. GDGT fluxes (μg·m^−2^·d^−1^) were normalized to deployment time and trap area.

### 2.5. Proxy calculations

The GDGT-based proxies used in this study are listed in Table 1. Fractional abundances of isoGDGTs were calculated relative to the total six isoGDGTs, and brGDGTs were calculated as fractions of the 15 brGDGTs. The following proxies were calculated: TEX_86_ (<u>Schouten et al., 2002</u>), branched and isoprenoid tetraether (BIT) (<u>Hopmans et al., 2004</u>), GDGT-0 to crenarchaeol ratio (GDGT-0/Cren) (<u>Blaga et al., 2009</u>), Ring Index (RI) (<u>Zhang et al., 2016a</u>), and crenarchaeol to crenarchaeol’ ratio (Cren/Cren’) (<u>Li et al., 2016</u>). For brGDGTs, the CBT was calculated according to <u>Weijers et al. (2007)</u>, and the degree of methylation of branched tetraethers of 5-methyl and 6-methyl brGDGTs (MBT’_5ME_, MBT’_6ME_), as well as Index 1 and the isomer ratio of 6-methyl brGDGTs, were determined following <u>De Jonge et al.</u> (<u>2014a)</u>; (<u>2014b</u>).

### 2.6. Statistical analysis

The relationship between water temperature and GDGT distributions, including both the fractional abundances of individual GDGTs and GDGT-based proxies, was evaluated using Pearson correlation analysis (Tables 1 and S1). As an initial step, we assessed the correlation between temperature and all GDGT-based proxies, regardless of whether they were originally designed as temperature proxies, in order to identify the most responsive compounds and proxies. Only those proxies that showed consistently strong and significant correlations with water temperature were selected for further discussion in this study. Water temperature was characterized using three metrics with different temporal scales: the daily mean temperature (DMT), the multiple-day mean temperature (MMT), and the mean temperature of each sampling interval. The DMT was defined as the average water temperature for a single day. In this study, DMT values were compiled for the mean temperature on a specific day within the 60 days prior to the trap recovery date, representing the long-term background thermal conditions. DMT_0_ represents the daily mean temperature on the trap recovery date, while DMT_–60_ corresponds to the temperature 60 days before trap recovery.

To explore the sensitivity of GDGTs to temperature across varying temporal scales, we employed a moving-window averaging approach to calculate MMT based on DMT, using window sizes ranging from 1 to 30 days (Fig. S1a). When the window size is 1 day, MMT_1_ is identical to DMT, and for larger window sizes, MMT is calculated over multiple overlapping intervals within the 60-day pre-recovery period. For example, with a 20-day window, 42 overlapping time windows (Window_0_ to Window_41_) can be defined, Window_0_ spans from DMT_–19_ to DMT_0_, Window_1_ corresponds to DMT_–20_ to DMT_1_, and so forth, with Window_41_ covering DMT_–60_ to DMT_–41_ (Fig. S1b). This systematic moving-window approach enables identification of the temperature segment most relevant to GDGT signals, without presupposing whether the beginning, middle, or end of the deployment period is most representative. The 60-day range includes windows centered near the midpoint, addressing concerns about temporal alignment. Among all tested windows, the 20-day interval (MMT_20_) showed the most consistent and robust correlation with GDGTs across various window sizes (see Discussion 4.2). Therefore, MMT_20_ was selected for subsequent analyses of temporal lag and vertical offset in GDGT–temperature relationships.

To further evaluate the vertical coherence of these relationships, MMT_20_ values were calculated at high vertical resolution, 0.5 intervals from 0 to 20 m, and 2 m intervals from 20 to 40 m, to assess spatiotemporal coupling between temperature profile and GDGT distributions in the water column.

All statistical analyses were performed in R (version 4.3.2). Pearson correlations were computed using the cor.test () function from the stats package. MMT values were calculated using the movavg () function from the pracma package (version 2.4.4).

## 3. Results

### 3.1. Temperature profile of the water column

The 19-month water temperature monitoring revealed that no ice formation occurred during the winter of 2012–2013. Throughout the monitoring period, the water temperature at 40 m depth remained relatively stable at approximately 8 °C (Fig. 1b), consistent with previous observations (<u>Wen et al., 2016</u>). Temperature variations were more pronounced in the upper water column compared to the deeper layers. For instance, at 10 m depth, the temperature ranged over 10 °C, while at 35 m, it was less than 2 °C (Fig. 1b). The temporal temperature profile confirmed the monomictic nature of Lake Lugu: the upper water column became thermally stratified from May to October, followed by cooling and full mixing from January to March. During winter mixing, vertical temperature differences were minimal (∼1.5 °C), whereas during strong stratification, temperature differences varied significantly with depth, decreasing with increasing depth (Fig. 1c).

### 3.2. Temporal distribution of GDGTs in settling particles

GDGTs were detected in all sediment trap samples (*n* = 32, Fig. 2). The mass flux of settling particles displayed a clear temporal pattern, with elevated fluxes during winter mixing and lower fluxes during stratified conditions (Fig. 2). Similarly, both the summed and individual GDGT fluxes followed this same temporal trend, peaking during late winter to early spring mixing, and reaching the minimum during summer stratification (Fig. 2). Throughout the sampling period, the flux of summed brGDGTs (∑brGDGTs) was consistently higher than that of summed isoGDGTs (∑isoGDGTs) (Fig. 2). A strong correlation was observed between ∑brGDGTs flux and total mass flux (R^2^ = 0.74, *p* < 0.05), while the correlation for ∑isoGDGTs was weaker (R^2^ = 0.47, *p* < 0.05) (Fig. S2). The ∑isoGDGTs fluxes spanned three orders of magnitude, ranging from 0.02 μg·m^−2^·d^−1^ in June 2013 to 21.7 μg·m^−2^·d^−1^ in January 2014. In contrast, ∑brGDGTs fluxes exhibited less variability, ranging from 0.06 to 4.1 μg·m^−2^·d^−1^ over the same period (Fig. 2 and Table S2).

**Fig. 2.**
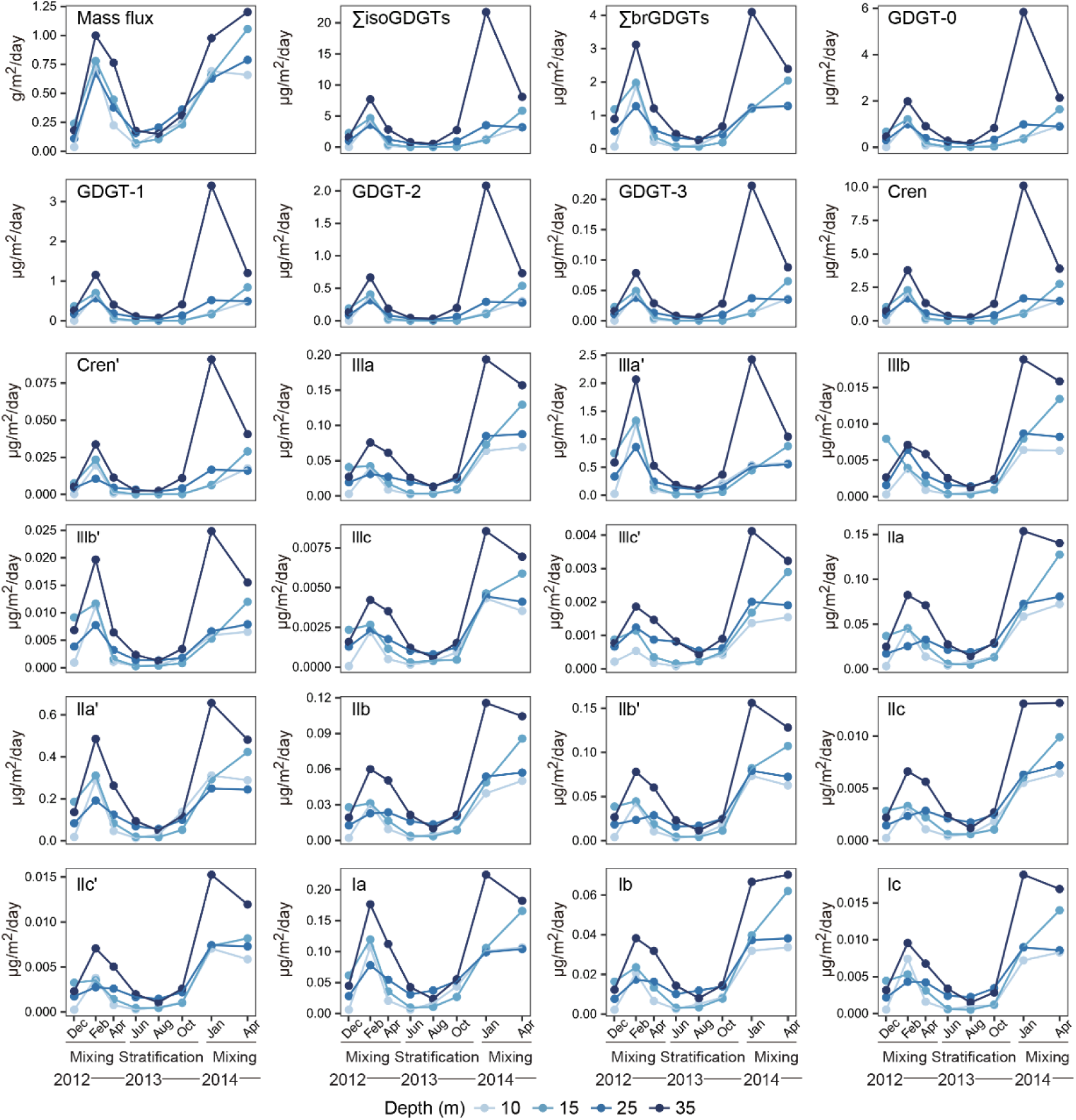
Temporal and spatial variation in mass flux (g·m^-2^·d^-1^), brGDGT, and isoGDGT fluxes (μg·m^-2^·d^-1^) of settling particles. Settling particles were collected from sediment traps deployed at four depths and recovered at approximately bimonthly or trimonthly intervals between December 2012 and April 2014. The periods of water column stratification and mixing are also indicated.

The isoGDGTs pool was dominated by GDGT-0 and crenarchaeol, which together accounted for up to 80% of the total isoGDGTs, respectively. GDGT-1, –2, –3 and Cren’ were minor components, contributing less than 20% (Figs. 3a and S3). Seasonal variation in the fractional abundances of isoGDGTs was evident: GDGT-0 was more abundant during stratification, especially at shallow depth, whereas crenarchaeol was enriched during mixing (Figs. 3c and e). In contrast, brGDGT compositions were relatively stable across seasons (Figs. 3b, d and f). The 6-methyl isomers of brGDGTs, IIIa’ and IIa’, were dominant, accounting for ∼60 and 30 % of the total brGDGTs pool, respectively, followed by the 5-methyl isomers Ia and IIa. The predominance of 6-methyl isomers resulted in consistently high IR_6ME_ values, averaging 0.83 ± 0.05 (Table S2).

**Fig. 3.**
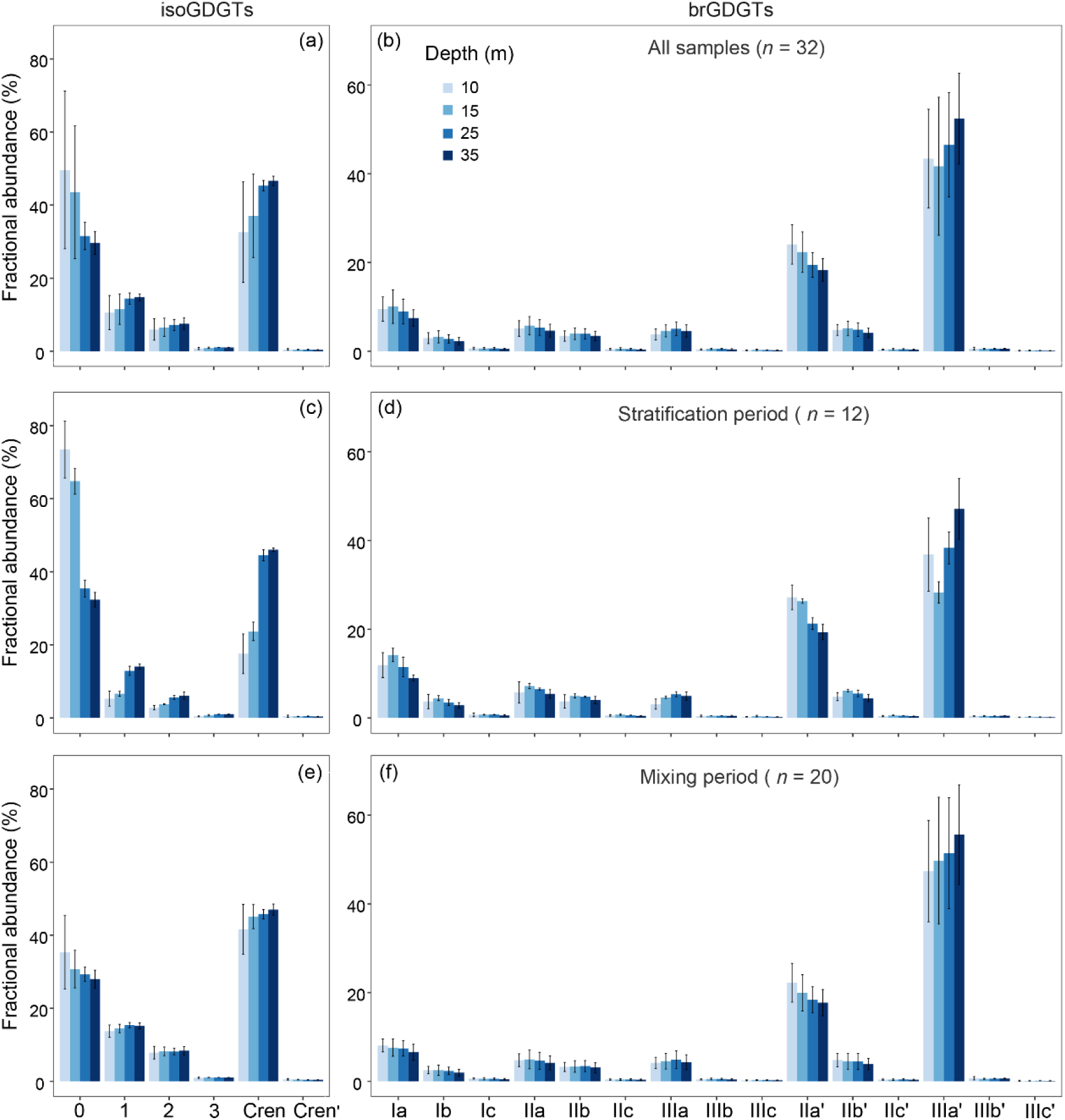
Average and standard deviation of the fractional abundance of isoGDGTs. (a) and brGDGTs (b) in settling particles. A total of 32 trap samples were collected from four depths across eight sampling intervals between December 2012 and April 2014. The fractional abundance of GDGTs during the periods of summer stratification (c and d) and winter mixing (e and f) is also shown.

These compositional variations significantly influenced GDGT-based proxies, particularly for traps deployed at the two shallow depths (10 and 15 m) (Fig. 4). The RI values varied from 0.6 to 2.3, generally showing lower values during summer stratification (Fig. 4c and Table S2). In contrast, the BIT and GDGT-0/Cren ratios were elevated under stratified conditions (Figs. 4b and d), with the highest GDGT-0/Cren ratio recorded in August 2013 due to high GDGT-0 and low crenarchaeol concentrations (Fig. S3). Periods of low Cren/Cren’ ratios coincided with high BIT and GDGT-0/Cren ratios (Figs. 4b, d and f). TEX_86_ values, however, did not show a clear seasonal pattern and fluctuated between 0.3 to 0.5 (Fig. 4a). Index 1 and MBT’_6ME_ were generally higher during stratification (Figs. 4g and i), while other brGDGT-based proxies such as CBT and MBT’_5ME_ showed limited temporal variability (Figs. 4e and h).

**Fig. 4.**
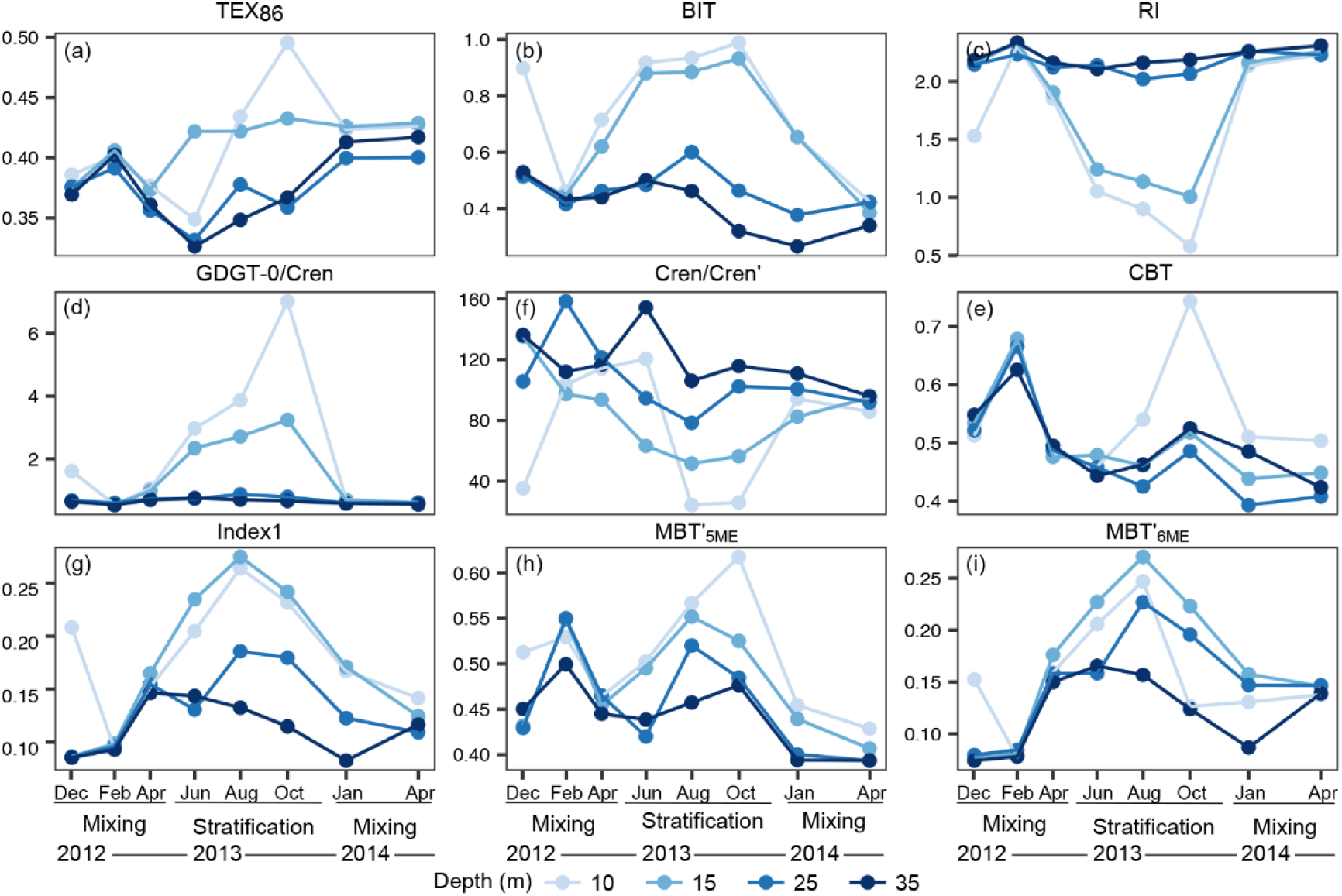
Temporal and spatial variations of GDGT-based proxies in settling particles collected from four depths between December 2012 and April 2014. The analyzed GDGT-based proxies include: (a) TEX_86_, (b) BIT, (c) RI, (d) GDGT-0/Cren, (e) Cren/Cren’, (f) CBT, (g) Index 1, (h) MBT’_5ME_, and (i) MBT’_6ME_. The calculations and definitions for these proxies are provided in Table 1.

### 3.3. Vertical distribution of GDGTs in settling particles

In addition to temporal variations, both GDGT fluxes and compositions varied with depth (Figs. 2–4). For analytical clarity, sediment traps were categorized by depth: 10 and 15 m (shallow), 25 m (intermediate), and 35 m (deep). The highest GDGT fluxes were observed at 35 m in January 2014, with the mean ∑isoGDGTs flux reaching 5.8 μg·m^−2^·d^−1^, significantly higher than at 10 m (1.1 μg·m^−2^·d^−1^) (Fig. 2 and Table S2). In contrast, no consistent increase was observed among the shallow depths. Rather, GDGT fluxes at 10 and 15 m were sometimes higher than at 25 m, such as during February 2013 (Fig. 2 and Table S2). Additionally, normalization to mass flux did not reveal a systematic enrichment of any GDGTs with increasing depth across this shallow interval.

At shallow depths, GDGT-0 was the dominant isoGDGTs, comprising 40–50 % of the total, but its relative abundance decreased to ∼30% at deeper depths (Fig. 3a). Conversely, crenarchaeol increased from ∼35% at shallow depths to over 45% at 35 m (Fig. 3a). These shifts resulted in higher GDGT-0/Cren ratio at shallow depths and low ratios at depth (Fig. 4d). Other cyclic isoGDGTs (GDGT-1, –2 and –3) also increased with depth (Fig. 3a).

IsoGDGT-based proxies reflected these vertical trends: RI increased from an average value of 1.7 at shallow depths to 2.2 at depth (Fig. 4c and Table S2). BIT values were higher at shallow depths (mean = 0.7) than at deeper depths (mean = 0.4) (Fig. 4b and Table S2). The mean TEX_86_ values ranged from 0.41 at shallow depths to ∼ 0.37 at depth. Overall, isoGDGT distributions and their proxies showed greater variability at shallow depths and were more stable in deep waters (Figs. 3a and 4).

The vertical pattern of ∑brGDGTs flux followed a similar trend to that of ∑isoGDGTs flux. It was lower at 10 m (mean = 0.7 μg·m^−2^·d^−1^) and increased to 1.6 μg·m^−2^·d^−1^ at 35 m (Fig. 2 and Table S2). However, unlike isoGDGTs, the fractional abundance of brGDGTs showed little variation with depth (Figs. 3b and S2). IIIa’ was the most abundant brGDGTs at 10 m (mean = 40% of the total brGDGTs), increasing to 50% at 35 m. Its flux also rose from 0.35 to 0.92 μg·m^−2^·d^−1^ with depth (Fig. 2 and Table S2). IIa’, the second-most abundant compound, accounted for 15–30% of brGDGTs at all depths (Figs. S3 and 3b). Its mean fractional abundance decreased slightly from 25% at 10 m to 20% at 35 m, while its flux increased from 0.14 to 0.29 μg·m^−2^·d^−1^ (Figs. 2, 3b and Table S2).

## 4. Discussion

### 4.1. Sources of GDGTs in the water column

In lacustrine environments, GDGTs preserved in sediments typically originate from a combination of terrestrial, aquatic, and sedimentary sources. To assess the origin of br– and isoGDGTs in the water column of Lake Lugu, we first examined the correlations between the bulk mass flux and the summed fluxes of br– and isoGDGTs, respectively (Fig. S2). The ∑brGDGTs flux exhibited a stronger correlation with mass flux (R^2^ = 0.74; Fig. S2b) than the ∑isoGDGTs flux (R^2^ = 0.47; Fig. S2a), suggesting that brGDGT production is more closely associated with overall particle production within the water column. Despite this difference, both ∑br– and ∑isoGDGTs fluxes followed similar temporal trends, increasing during winter mixing and decreasing during summer stratification. This co-variation resulted in a strong correlation between the fluxes of ∑br– and ∑isoGDGTs (R^2^ = 0.83; Fig. S2c), implying that the production of bacterial and archaeal GDGTs in settling particles may share a common autochthonous origin or be governed by similar environmental factors. Additionally, both summed and individual GDGT fluxes increased with the depth (Fig. 2), consistent with in situ microbial production in the water column rather than external delivery.

This conclusion is further reinforced by a combination of field data and prior studies. First, GDGT fluxes were consistently low during the rainy season, when terrestrial input would be expected to be maximum (Fig. 2). This inverse relationship suggests that terrestrial contributions were minimal. Second, the observed depth-dependent flux patterns were inconsistent with a surface-derived source, and instead imply active production within the lake. Third, previous study from Lake Lugu revealed substantial differences in isoGDGT distributions and concentrations between surface sediments and surrounding soils (<u>Li et al., 2023</u>), suggesting predominant in situ production within the lake. Similarly, depth-related increases in brGDGT concentrations across 54 surface sediment samples (Li et al., unpublished data) further support an autochthonous source of brGDGTs in this deep alpine lake. However, the pronounced increase in GDGT fluxes at 35 m warrants further investigation. Although part of this enhancement may be attributed to intensified in situ production in the hypolimnion, the magnitude of the increase, especially when compared to the relatively consistent fluxes between 10 and 25 m, suggests that physical processes, such as resuspension or vertical particle transport may also play a significant role.

While we acknowledge that lateral transport and shoreline processes may contribute to the observed signals, particularly at specific depths or time intervals, the consistent depth-related and seasonal patterns suggest that in situ microbial production is the dominant source of GDGTs in the water column.

These production patterns align with the known ecological niches of GDGT-producing microorganisms. In particular, the nitrifying Thaumarchaeota, primary producers of isoGDGTs, are known to thrive in deep, oxygenated waters with elevated ammonium concentrations (<u>Buckles et al., 2013</u>; <u>Baxter et al., 2021</u>; <u>Sinninghe Damsté et al., 2022</u>). In Lake Lugu, DO concentrations vary with season and depth but generally range from 3 to 8 mg l^−1^ in the upper 60 m (<u>Wang et al., 2018</u>). During winter mixing, DO levels are relatively constant at 8 mg l^−1^ throughout the water column, creating favorable conditions for Thaumarchaeotal activity. These observations are consistent with previous findings suggesting that Thaumarchaeota Group I.1a are outcompeted in the upper oxic layer (<u>Buckles et al., 2013</u>; <u>Kumar et al., 2019</u>; <u>Baxter et al., 2021</u>). Together, these data suggest that most isoGDGTs in settling particles originate from in situ production in the water column, with minimal contributions from terrestrial sources, particularly during the low-flux summer rainy season.

This interpretation is further supported by the elevated GDGT-0/Cren ratios (> 2) observed at the two shallow depths during the summer stratification period (Fig. 4d and Table S2). Since Thaumarchaeota typically produce high amounts of crenarchaeol but relatively little GDGT-0, GDGT-0/Cren ratio in pure Thaumarchaeota cultures are generally < 2 (<u>Blaga et al., 2009</u>; <u>Sinninghe Damsté et al., 2012b</u>; <u>Naeher et al., 2014</u>).

The higher ratios observed at shallow depths during stratification therefore likely reflect increased contribution from non-Thaumarchaeotal archaea. Additionally, the consistently high Cren/Cren’ ratios (> 25) throughout the study period suggest a low relative contribution of Group I.1b Thaumarchaeota to the isoGDGTs pool (<u>Sinninghe Damsté et al., 2012b</u>; <u>Li et al., 2016</u>) (Fig. 4f and Table S2). This inference is further supported by DNA sequencing data from Lake Lugu surface sediments, which indicate that *Nitrosoarchaeum* (affiliated with Group I.1a) is the dominant Thaumarchaeal operational taxonomic units (OTU) in this system (<u>Ren and Wang, 2022</u>).

### 4.2. Temporal lag between temperature and GDGTs

To investigate the temporal lag between water temperature and GDGT distributions, we compared in situ measured temperature (at trap deployment depths) with the GDGTs composition of settling particles collected from four different depths. Temperature data were categorized into four types based on temporal resolution: (1) the mean temperature over the entire study period, (2) the mean temperature for each sampling interval, (3) the multiple-day mean temperature (MMT) with various window sizes, and (4) the daily mean temperature (DMT). Since the full-period mean temperature is too coarse to resolve potential time-lagged responses, we focused on interval-averaged and higher-resolution daily and multi-day metrics in this analysis.

Preliminary correlation analyses using mean water temperatures for each sampling interval showed that isoGDGTs, particularly GDGT-0 and crenarchaeol, were more strongly correlated with temperature than brGDGTs (Fig. S4a). Among GDGT-based proxies, the RI exhibited the strongest temperature sensitivity, outperforming TEX_86_ (Fig. S4b). These initial results provided the rationale for investigating potential temporal lags using high-resolution daily and multiple-day temperature records (see Supplementary Fig. S4 for details).

#### 4.2.1. Relationship between daily mean temperature and GDGTs

While interval-averaged temperatures can reveal general seasonal patterns, they are insufficient for capturing short-term or time-lagged responses in GDGTs. To address this limitation, we calculated daily mean temperature (DMT) values over the 60 days prior to each trap recovery date, thereby standardizing the temporal resolution. DMT was determined by averaging in situ water temperatures recorded every two hours at each trap depth during the 60-day period (Fig. S5). Meta-analysis of the DMT–GDGT relationship revealed that isoGDGTs exhibit greater sensitivity to short-term temperature variations than brGDGTs, consistent with results based on interval-averaged temperatures (Fig. S4). For instance, the fractional abundance of GDGT-0 showed a strong positive correlation with DMT, while crenarchaeol was negatively correlated (Figs. S5a and b). Notably, both compounds exhibited peak correlations approximately 20 days prior to trap recovery (DMT_–20_), with absolute Pearson correlation coefficients (|*r*|) reaching 0.93 (*p* < 0.001). Similarly, the RI displayed a strong negative correlation with DMT, also peaking at DMT_–20_ (|*r*| = 0.93; Fig. S5c).

These results highlight the strong thermal dependence of isoGDGTs and further support their use as sensitive proxies for past lake water temperatures. As illustrated in the circos plots, correlation patterns over the entire 60-day window confirm this ∼20-day temporal lag: correlation coefficients increase from the date of trap recovery up to ∼20 days prior (*r*_max_= 0.93; Figs. S6b and d), and then gradually decline toward day –60 (Figs. S6a and c). Additional isoGDGT compounds, such as GDGT-1 and GDGT-3 showed similar peak correlations around the 20-day mark (Figs. S6e and S7), while BIT and GDGT-0/Cren ratios also exhibited maximum correlations at DMT_–20_ (Figs. S6f and S7).

Interestingly, RI consistently exhibited a stronger correlation with temperature than TEX_86_ (Figs. S4b and S6f), likely due to its dependence on the relative abundances of GDGT-0 and crenarchaeol, which are sensitive to thermal variations. This finding aligns with culture-based studies of Thaumarchaeota, where RI displayed stronger correlations with temperature than TEX_86_ (R^2^ = 0.86 vs. 0.56; Fig. S8) (<u>Elling et al., 2015</u>; <u>Qin et al., 2015</u>; <u>Zhang et al., 2016a</u>; <u>Elling et al., 2017</u>). However, this contrasts with findings from marine mesocosm experiments and core-top sediments, where both RI and TEX_86_ typically co-vary with temperature (<u>Elling et al., 2015</u>; <u>Zhang et al., 2016a</u>). In our study, as well as in previous investigations of surface sediments, suspended particulate matter, and settling particles from non-marine environments, no significant correlation has been observed between these two proxies (Fig. S9). This broader inconsistency is further supported by global peatland datasets, where diverse archaeal communities are thought to obscure the temperature dependence of isoGDGTs (<u>Naafs et al., 2018</u>).

In contrast to isoGDGTs, brGDGTs showed limited correlation with temperature in Lake Lugu. This includes the MBT’_5ME_, which despite performing well in global lacustrine calibration datasets (<u>Russell et al., 2018</u>; <u>Martínez-Sosa et al., 2021</u>; <u>Raberg et al., 2021</u>), as well as in mineral soils and peats (<u>De Jonge et al., 2014a</u>; <u>Naafs et al., 2017b</u>), did not correlate strongly with temperature in settling particles. The relatively high abundance of hexamethylated brGDGTs and the low proportion of tetramethylated brGDGTs observed in our samples conform to the brGDGT distribution patterns observed in global lake surface sediments (Fig. S10), suggesting a broadly consist source signal. However, despite the brGDGT-based paleothermometers have demonstrated strong correlations with temperature in these surface sediment datasets (<u>Martínez-Sosa et al., 2021</u>; <u>Raberg et al., 2022</u>), such relationships were not evident in the settling particles examined in this study.

This discrepancy suggests that environmental factors other than temperature, particularly DO and nutrient availability, may exert a stronger influence on brGDGT production in the water column. For instance, spatial variations in bacterial communities across the oxycline have been shown to shape brGDGT distributions independently of temperature in Lake Lugano, a deep meromictic lake in Switzerland (<u>Weber et al., 2018</u>). Similarly, in Lake Chala, water column stratification dynamics rather than temperature, appear to be the main driver of brGDGTs production (<u>vanBree et al., 2020</u>). In a Chinese lake, DO levels have been found to significantly affect brGDGT distributions in surface sediments (<u>Wu et al., 2021</u>). More recently, a pure culture study of *Edaphobacter aggregans*, a known brGDGTs-producing *Acidobacterium*, demonstrated that oxygen availability directly regulates brGDGTs production (<u>Halamka et al., 2021</u>). Collectively, these findings support the hypothesis that oxygen concentration, rather than temperature, may be the primary driver of brGDGTs production in certain lake environments, including Lake Lugu. We recommend that future studies incorporating settling particles account for DO dynamics and other environmental variables when interpreting brGDGT-based paleoclimate proxies.

#### 4.2.2. Relationship between the multiple-day mean temperature and GDGTs

To further evaluate the ∼20-day temporal lag observed between DMT and GDGT distributions, we examined the relationship between the multiple-day mean temperature (MMT) and GDGTs. This analysis aims to quantify the integrated thermal signal preserved in settling particles and to complement the DMT-based correlation analysis by capturing time-averaged effects over varying temporal scales. MMT values were determined using a moving-window averaging approach (<u>Kristoufek, 2014</u>; <u>Felipe-Lucia et al., 2020</u>) applied to the DMT time series, with window sizes ranging from 1 to 30 days prior to each trap recovery (Fig. S1a). As the window size increases, the number of available windows decreases accordingly. This framework allows for the assessment of GDGT–temperature correlations across both short– and intermediate-term time scales.

To identify the optimal integration period, we selected the RI, the most temperature-sensitive GDGT-based proxy in our dataset, as a representative indicator. Pearson correlation coefficients between RI and MMT were calculated for each combination of window length and position., RI consistently exhibited significant negative correlations with MMT across all window sizes, with the strongest correlation observed at MMT_20_ (highlighted in red in Fig. S11), confirming its robust thermal sensitivity. Shorter windows (<10 days) produced stronger correlation magnitudes but higher noise due to transient fluctuations, while longer windows (>25 days) yielded smoother trends at the expense of correlation strength due to signal averaging (Fig. S11). Considering these trade-offs, the 20-day MMT (MMT_20_) was selected as optimal for subsequent analyses, balancing signal robustness with sensitivity to temperature variability.

Circos plots and corresponding scatter plots depicting the temporal evolution of Pearson correlations between MMT_20_ and GDGTs further confirmed the observed ∼20-day thermal integration window (Fig. 5). GDGT-0 and RI, the most temperature-sensitive GDGT components, showed peak correlations with MMT_20_ centered around the trap recovery date, with *r* values reaching 0.93 (Figs. 5a–d). Correlation strength gradually declined as the window lag increased beyond 20 days. Similar but slightly weaker trends were observed for other isoGDGTs and proxies, including GDGT-1, GDGT-3, BIT, and GDGT-0/Cren (Figs. 5e and f). These consistent patterns indicate that GDGT signals in settling particles predominantly reflect integrated thermal conditions over the ∼20 days preceding collection.

**Fig. 5.**
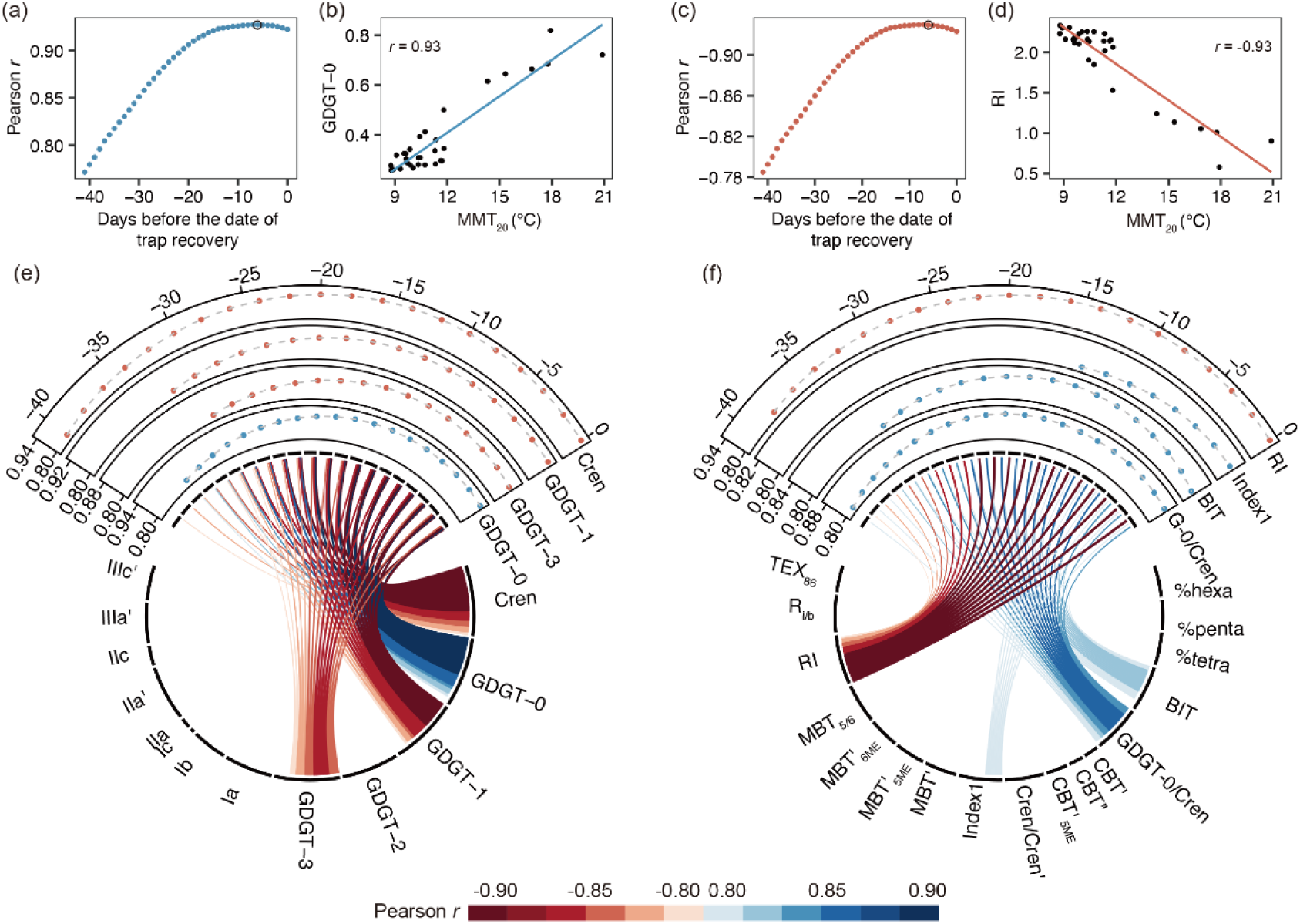
Pearson correlation analysis between the multiple-day mean temperature with a 20-day window (MMT_20_) and GDGTs. GDGTs are expressed by their fractional abundances and GDGT-based proxies. Panels (a) and (c) show the variation in Pearson correlation coefficients (*r*) between MMT_20_ calculated over 42 overlapping windows prior to trap recovery and the fractional abundance of GDGT-0 and the RI, respectively. The maximum absolute correlation values (|*r*|) for GDGT-0 and RI are highlighted by black circles. Panels (b) and (d) present scatter plots illustrating the relationships between MMT_20_ and the fractional abundances of GDGT-0 and RI, respectively. Circos plots (e) and (f) visualize the variation in *r* between MMT_20_ and the fractional abundances of individual GDGTs and GDGT-based proxies, respectively. In the circos plot, the upper quarter highlights the four variables with the strongest correlations, with Pearson |*r*| > 0.8. Along the top arc, 21 data points represent averaged Pearson correlation coefficients from pairs of adjacent MMT_20_ windows (out of 42), showing their relationships with the fractional abundances of GDGTs and GDGT-based proxies. Proxy definitions are provided in Tables 1 and S1.

Importantly, such a clearly ∼20-day temporal lag between water temperature and GDGTs in settling particles has rarely been quantified in lacustrine studies. While previous investigations have suggested offsets between GDGTs-reconstructed and instrumental temperatures in tropical and temperate lakes (<u>Buckles et al., 2014</u>; <u>Loomis et al., 2014</u>), the precise duration and mechanisms of these lags remain poorly constrained. The biological basis for the observed ∼20-day temporal lag remains uncertain. One possible explanation is partial in situ production of GDGTs within the sediment traps by microbial communities associated with settling or retained particles. These microbes may contribute newly synthesized GDGTs with intrinsic turnover times on the order of 20–30 days. However, this interpretation should be treated with caution, as GDGT turnover rates remain poorly constrained, and microbial community dynamics are known to vary widely in aquatic environments (<u>Spohn et al., 2016</u>; <u>Kritzberg and Bååth, 2022</u>).

Alternatively, GDGTs produced earlier during the trap deployment period may have undergone degradation or lateral export, thereby weakening correlations with earlier temperature signals. In addition to biological factors, physical processes may also play a role. For instance, resuspension in the early stage of trap deployment could introduce reworked or allochthonous material, diluting the contemporaneous temperature signal. As trap accumulation proceeds and the influence of resuspension declines, later-settling GDGTs may better reflect prevailing thermal conditions.

Taken together, these factors provide a plausible explanation for the strongest GDGT–temperature relationships emerging from the MMT_20_. They emphasize the importance of considering both microbial turnover and early diagenetic or physical processes when interpreting GDGT signals from sediment traps. The lack of similar observations in previous studies likely stems from the absence of high-resolution and continuous temperature data. Future investigations integrating fine-scale temperature monitoring, trap segmentation, and microbial community analysis would help to better constrain the drivers of temporal lags in GDGT production and preservation in lacustrine systems.

### 4.3. Spatiotemporal shift between temperature and GDGTs

To investigate the spatiotemporal offsets between GDGT signals and ambient water temperature, we performed depth-resolved correlation analyses using high-resolution temperature profiles (0–40 m) and GDGT distributions from four sediment trap depths (10, 15, 25, and 35 m). Among all GDGT-based proxies, the RI showed the strongest and most consistent temperature sensitivity, with a clear ∼20-day temporal lag in response to both daily mean temperature (DMT_–20_) and multiple-day mean temperature (MMT_20_) (Figs. 5 and S6).

To assess spatial offsets, RI values were correlated with MMT_20_ profiles resolved at 0.5 m intervals (0–20 m) and 2 m intervals (20–40 m) across 42 overlapping windows (Window_0_ to Window_41_). Each window represents a 20-day moving average of DMT starting from 0 to 41 days prior to trap recovery (see Section 2.6 and Fig. S1b). The correlation analysis revealed that RI at all depths was more strongly associated with MMT_20_ from shallower layers than from the deployment depth, indicating a consistent upward spatial offset in thermal signal captured by settling particles (Fig. 6). For instance, RI at 25 m depth exhibited its highest correlation with MMT_20_ at 4 m during Window_10_ (DMT_–29_ to DMT_–10_), with an *r* of –0.83, significantly higher than the correlation at its own depth (*r* = –0.59; Fig. 6b). This trend was more pronounced at 35 m, where the RI–MMT_20_ correlation was notably weak (*r* = –0.66 vs. –0.43 at 4 and 35 m; Fig. 6b), suggesting signal degradation at greater depths likely due to sediment resuspension.

**Fig. 6.**
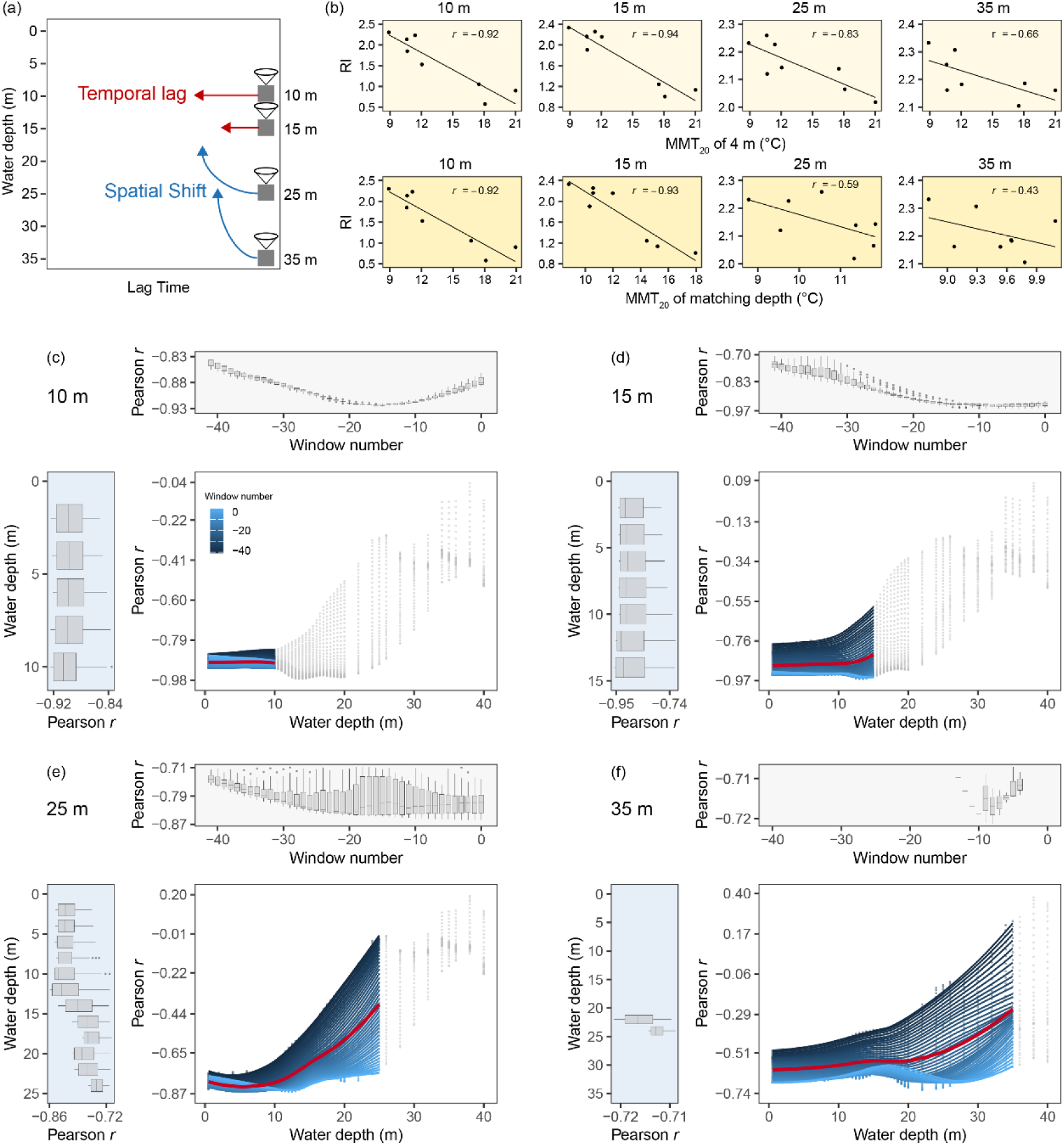
Temporal lags and vertical offsets in the relationship between MMT_20_ and the RI. The RI was calculated from isoGDGTs in settling particles collected at four depths (10, 15, 25 and 35 m). Panel (b) indicates the Pearson correlation coefficients between RI at each trap depth and MMT_20_ values calculated at 4 m (upper) and at the corresponding trap depth (lower). Panels (c–f) illustrate the spatiotemporal patterns of RI–MMT_20_ correlations for each trap depth, with each panel comprising three subplots: **Upper subpanels** show the temporal evolution of Pearson *r* between RI at each trap depth and MMT_20_ values at various depths above the trap, calculated across 42 overlapping time windows prior to trap recovery. Each box represents the distribution of correlation coefficients across all depths above the trap for a single time window, with the x-axis denoting the 42 sequential windows. **Left subpanels** display vertical correlation profiles between RI at each trap depth and MMT_20_ values at specific depths above the trap. Each box displays the distribution of the 42 correlation coefficients calculated over all time windows at a given depth, with the y-axis indicating water depth. **Central subpanels** integrate both temporal and spatial dimensions, combining depth-wise correlations between RI and MMT_20_ across all depths above the trap over the 42 overlapping time windows (blue shades). Gray points indicate correlation coefficients for depths below the trap. The fitted red curves represent loess-smoothed trends of depth-wise correlations. MMT_20_ values were calculated at 0.5 m intervals between 0 and 20 m, and at 2 m intervals between 20 and 40 m.

To further explore spatiotemporal decoupling, RI was regressed against MMT_20_ from different depths across all 42 overlapping windows (Fig. 6c–f). Depth-specific optimal windows varied (Fig. 6c–f, top panels): At 10 m, the strongest correlations occurred between Window_10_ and Window_20_, peaking at Window_15_ (DMT_–34_ to DMT_– 15_), with *r* = –0. 92 (Fig. 6c). At 15 m, the best fit was found near Window_5_ (*r* = –0. 96; Fig. 6d). At 25 m, peak correlations occurred in windows centered around 10–20 days before trap recovery (Fig. 6e), although the specific optimal window needs to be confirmed. At 35 m, correlations remained weak regardless of windows, peaking at Window_10_ with of *r* = –0. 72 (Fig. 6f).

Additionally, the correlation between RI and MMT_20_ also exhibited clear depth-dependent spatial offsets (Fig. 6c–f, left panels). At 10 m, the strongest correlation was observed at the same depth (*r* = –0.93; Fig. 6c), indicating minimal spatial bias. At 15 m, the highest correlation occurred at 13 m (*r* = –0.96; Fig. 6d), suggesting a minor upward shift of 2 m. At 25 m, the peak correlation was observed at 10 m (*r* = – 0.83; Fig. 6e), implying a ∼15 m upward offset. At 35 m, the highest correlation was observed at 20 m (*r* = –0.72; Fig. 6f), also reflecting an upward offset of ∼15 m, but with attenuated signal strength compared to shallower depths. This reduction in fidelity likely results from intensified vertical mixing and near-bottom sediment resuspension, which may dilute or obscure depth-specific temperature signals. These patterns suggest that isoGDGTs collected at 35 m may include contributions from upper water layers via particle settling, leading to a vertically integrated rather than locally produced signal. To our knowledge, this study represents the first quantitative assessment of vertical offsets in GDGT–temperature relationships within a stratified lacustrine water column.

Notably, brGDGTs, exemplified by the fractional abundance of Ia, exhibited different spatial behaviors (Figs. S12a–d, left panels). At 10 m, Ia correlated most strongly with MMT_20_ at the same depth (*r* = 0.89; Fig. S12a), indicating no vertical offset. At 15 m, the strongest correlation was at 10 m (*r* = 0.92; Fig. S12b), and at 25 m, it was at 12 m (*r* = 0.86; Fig. S12c), suggesting spatial shifts of ∼5 and 13 m, respectively. These findings imply that brGDGTs (e.g., Ia) may be more responsive to vertical transport or spatial gradients than short-term temporal variability, unlike the isoGDGT-based RI which consistently reflected a ∼20-day lag. The divergent behaviors of iso– and brGDGTs likely stem from distinct source origins: isoGDGTs primarily originate in the euphotic zone and are delivered via settling particles, while brGDGTs may be produced by benthic or near-bottom microbes less exposed to upper water column dynamics. This highlights the necessity for proxy-specific calibrations when reconstructing vertical thermal profiles from sediment trap GDGTs.

A comprehensive spatiotemporal correlation analysis further revealed depth-dependent decoupling patterns (Fig. 7). At 10 m, the best match occurred at the same depth with a ∼15-day lag (Fig. 7a). At 15 m, RI was most strongly correlated with MMT_20_ from 13 m, with a lag of 0–10 days (Fig. 7b). At 25 m, the strongest correlation was with MMT_20_ at 5–10 m, lagging by 0–20 days (Fig. 7c). However, at 35 m, no robust spatiotemporal correspondence was observed (Fig. 7d). These results suggest that GDGT–temperature coupling weakens with depth due to physical mixing and resuspension, especially near the lake bottom. Overall, the distinct responses across depths highlight the complex interplay between in situ microbial production, vertical transport, and thermal stratification in shaping GDGT-based proxies in lake systems.

**Fig. 7.**
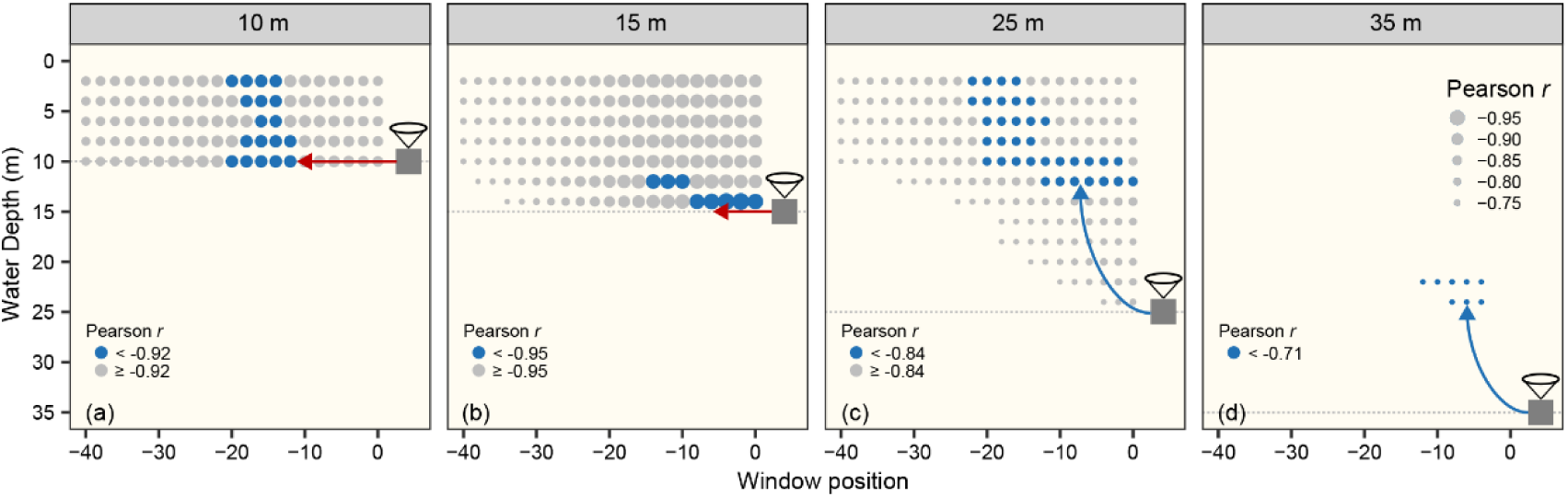
Conceptual model illustrating the temporal lag and vertical offset in the relationship between the RI of isoGDGTs from settling particles and MMT_20_. The model depicts distinct depth-dependent patterns in the RI–MMT_20_ correlations. At shallow depths (10 and 15 m; a and b), minimal vertical offsets were observed, indicating that the RI predominantly reflected the MMT_20_ at the same depth. At the intermediate depth (25 m; c), the RI exhibited strong correlations with MMT_20_ at shallow depths (10–15 m), implying downward transport of particles that had integrated surface-layer temperature signals with a temporal lag. At 35 m (d), both temporal and vertical correlations were comparatively weak, likely due to signal overprinting by near-bottom resuspension processes. Dot size represents the magnitude of the Pearson correlation coefficients between RI and MMT_20_, with larger dots indicating stronger correlations.

## 5. Conclusions

This study presents the first systematic assessment of the relationship between GDGTs in settling particles and in situ water column temperature of Lake Lugu, a deep, stratified alpine lake in southwestern China. Synchronous spatiotemporal variations in the fluxes of both brGDGTs and isoGDGTs across four depths provide evidence for in situ production within the water column, and possibly within the sediment traps as well. Among the isoGDGT-based proxies, the RI displays the strongest and most consistent correlation with temperature, albeit with an apparent ∼20-day temporal lag relative to contemporaneous water temperatures. This finding quantitatively confirms previously hypothesized but unverified time lags in GDGT– temperature coupling in lacustrine settings. Spatially, GDGT–temperature correlations reveal clear vertical offsets. Sediment traps at 10 and 15 m show minimal vertical offsets (0–2 m), suggesting that GDGTs at these depths primarily reflect local water thermal conditions. At 25 m, a pronounced upward offset of ∼15–20 m is observed, indicating that settling particles at this depth preferentially integrate temperature signals from the overlying water column. In contrast, the absence of a clear vertical correlation at 35 m is likely due to near-bottom sediment resuspension processes near the lake bottom (∼5 m above the sediment–water interface), which disrupt the vertical integrity of GDGT signals. Notably, brGDGT-based proxy (MBT’_5ME_) shows no significant relationship with temperature across any depth or time interval, contrasting with the temperature-sensitivity of the isoGDGT-based RI. This divergence likely reflects differences in the ecological origins of the two GDGT producers, with brGDGTs likely sourced from benthic or near-sediment microbial communities that are less influenced by water column thermal gradients.

By quantifying both temporal (∼20 days) and spatial (∼15–20 m) offsets in GDGT– temperature relationships, this study provides new mechanistic insight into how particle settling dynamics and microbial ecology shape GDGT signals in stratified lakes. These findings challenge the assumption that GDGTs directly reflect local temperature conditions, and call for greater consideration of vertical transport and in situ production processes in proxy calibration. Future research should prioritize multi-site comparisons, sediment trap–core integration, and compound-specific isotope analyses to further constrain GDGT production and export pathways. By addressing the spatiotemporal complexity of GDGT behavior, this work enhances the accuracy and resolution of GDGT-based paleoclimate reconstructions in alpine stratified lake systems.

## Supporting information

main text table

## Acknowledgements

This study was supported by the National Natural Science Foundation of China (Grant Nos. 42330511, 41977384, and 42225708), J.L. was funded by Science and Technology Planning Project of Nanjing Institute of Geography and Limnology, Chinese Academy of Sciences (NIGLAS2022TJ04). R.W. acknowledges the financial supports of the Youth Scientists Group in Nanjing Institute of Geography and Limnology, Chinese Academy of Sciences (2021NIGLAS-CJH03). B.D.A.N. acknowledges a Royal Society Tata University Research Fellowship for funding. The data used are listed in the tables, figures, and supplementary material of the paper. We deeply appreciate the Executive Editor and the associate editors for their patience and understanding during the extended revision period, particularly given the first author’s maternity leave. We extend our gratitude to the three anonymous reviewers for their detailed and insightful feedback which substantially improved this manuscript.

## Appendix A. Supplementary material

This supplementary file contains 12 figures, 2 tables, and corresponding references. Fig. S1: Schematic overview of MMT and window calculation. Fig. S2: Scatter plots showing relationships between mass flux and GDGT fluxes. Fig. S3: Temporal and spatial variation in the fractional abundance of GDGTs. Fig. S4: Pearson correlation matrices between mean water temperature during each sampling interval and GDGTs. Fig. S5: Correlation analysis between daily mean temperature (DMT) and GDGT components. Fig. S6: Circos plots illustrating Pearson correlation between DMT and GDGTs. Fig. S7: Scatter plots presenting correlation analyses between DMT and GDGTs. Fig. S8: Relationships between incubation temperature and TEX_86_ and RI from Thaumarchaeota culture experiments. Fig. S9: Comparisons between RI and TEX_86_ in sedimentary archives and particulate samples. Fig. S10: Ternary diagram of brGDGTs from this study and global lake surface sediments. Fig. S11: Variability in Pearson correlation coefficients between RI and MMT across different window sizes. Fig. S12: Temporal lags and vertical offsets in the relationship between MMT_20_ and Ia. Table S1: Calculation details for GDGT-based proxies. Table S2: Fluxes and GDGT-based proxies of all samples.

