## Supplemental materials for "Quantifying spatiotemporal decoupling of GDGT-temperature relationships in a deep alpine lake"

Contents:

-Supplementary Figures S1-12

-Supplementary Tables S1-2

Related references

### Figures

**Fig. S1.** Calculations of multiple-day mean temperature (MMT) and sliding window method. (a) The multiple-day mean temperature (MMT) was calculated using sliding windows based on the DMT values, with window sizes ranging from 1 to 30 days. For instance, when the window size is 1 day ($MMT_1$), the value corresponds directly to the DMT, yielding 61 overlapping windows over the 60-day pre-recovery period. A 2-day window produces 60 overlapping windows, a 20-day window generates 42 windows, and a 30-day window results in 32 windows. Each window represents a specific *n*-day averaging interval (window size) ending at a defined number of days (window position) prior to trap recovery. These windows are denoted as $DMT_{-(n-1)}$ to $DMT_0$ ($Window_0$), $DMT_{-n}$ to $DMT_{-1}$ ($Window_1$), ..., and $DMT_{-60}$ to $DMT_{-(61-n)}$ ($Window_m$), where $n$ is the window size and $m = 61 - n$. A 20-day window ($MMT_{20}$) was selected for subsequent analysis due to its stable and representative performance across various window sizes. (b) To assess potential time-lag and vertical effects between GDGT distributions and water temperature, we generated 42 overlapping $MMT_{20}$ windows, spanning from day –60 to day 0 relative to trap recovery. $Window_0$ corresponds to $DMT_{-19}$ to $DMT_0$, $Window_1$ spans from $DMT_{-20}$ to $DMT_{-1}$, and so on, with $Window_{41}$ covering $DMT_{-60}$ to $DMT_{-41}$, representing the thermal history prior to deposition. Pearson correlation coefficients were calculated between GDGT distributions and $MMT_{20}$ at 0.5 m intervals from 0 to 20 m, and at 2 m intervals from 20 to 40 m depth. Day 0 represents the trap recovery date, while day –60 refers to 60 days before trap recovery.

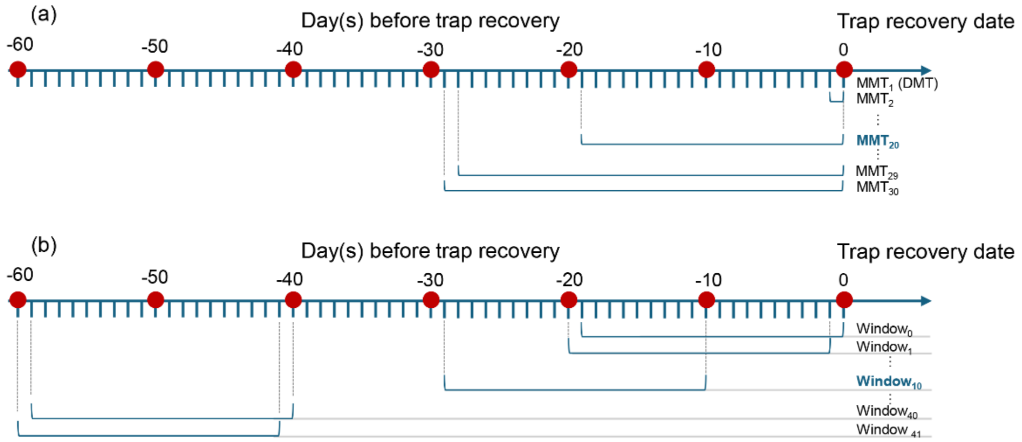

**Fig. S2.** Relationship between mass flux and GDGT fluxes in settling particles from Lake Lugu. (a) Scatter plots illustrating the relationship between mass flux and the ΣisoGDGTs flux; (b) the relationship between mass flux and the ΣbrGDGTs flux; (c) the relationship between the fluxes of ΣisoGDGTs and ΣbrGDGTs. Settling particles were collected using sediment traps deployed at four depths and retrieved at approximately bimonthly or trimonthly intervals from December 2012 to April 2014.

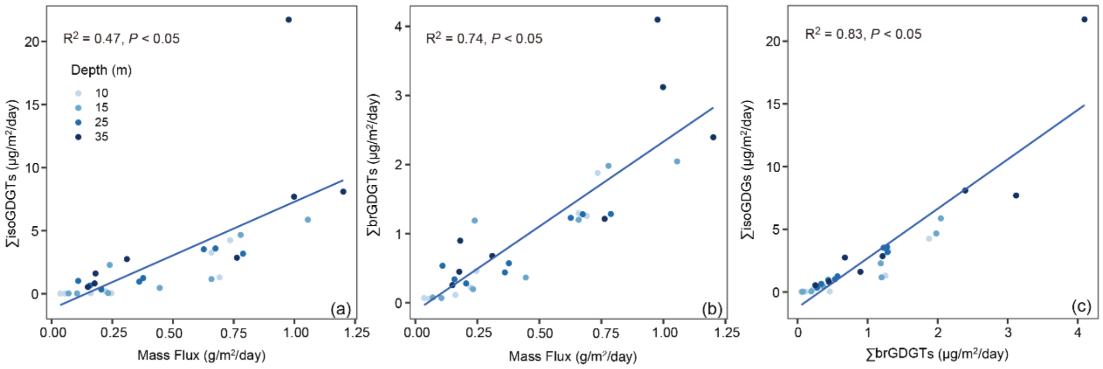

**Fig. S3.** Temporal and spatial variation in the fractional abundance of individual br- and isoGDGTs in settling particles from Lake Lugu. Settling particles were collected from sediment traps deployed at four water depths and recovered at approximately bimonthly or trimonthly intervals between December 2012 and April 2014.

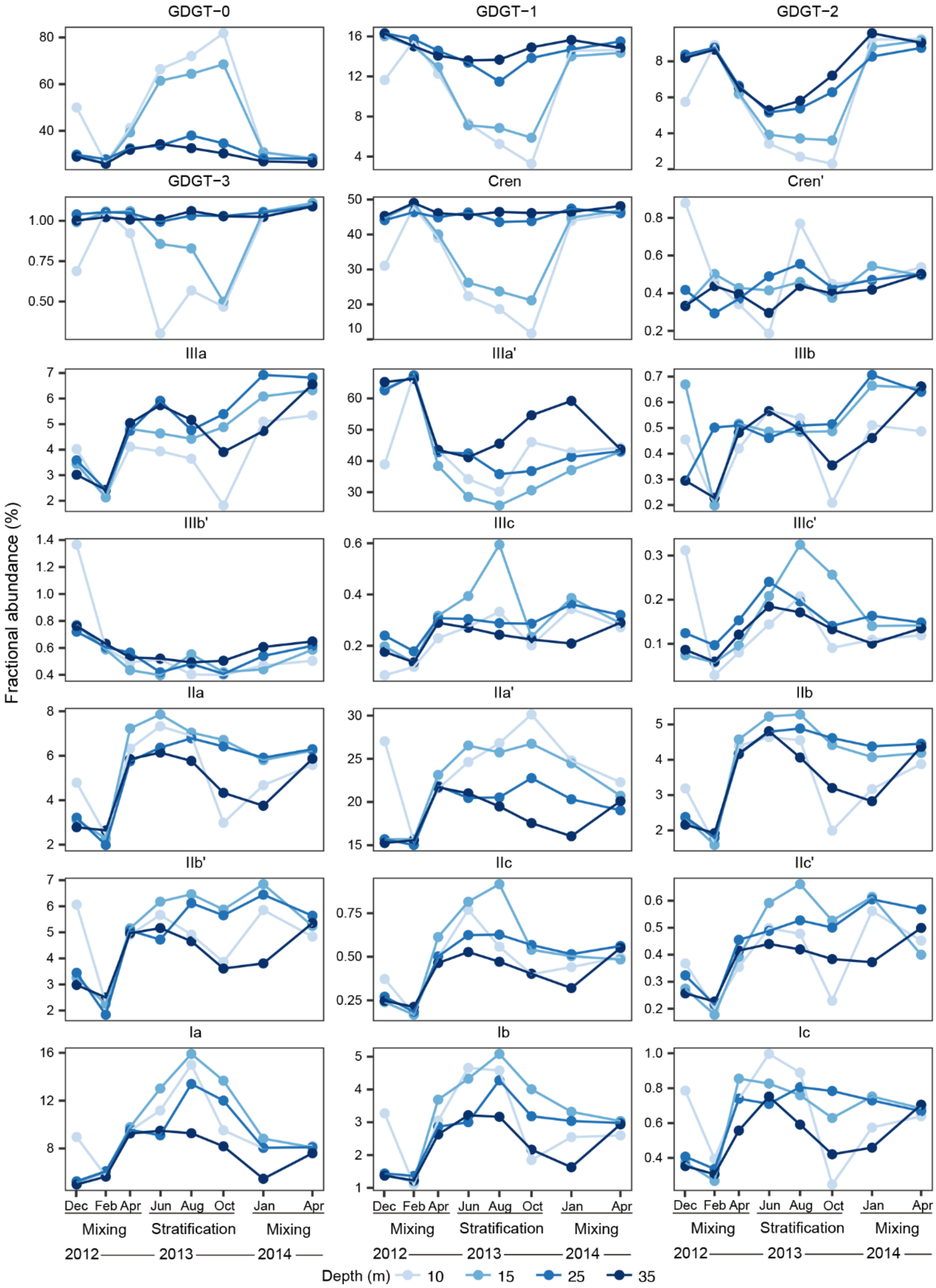

**Fig. S4.** Pearson correlation matrices between the mean water temperature for each sampling interval and GDGT distributions in settling particles. (a) Correlation coefficients (*r*) between the fractional abundances of individual GDGTs and the mean water temperature for each sampling interval. The analysis reveals that isoGDGTs exhibit a stronger sensitivity to temperature than brGDGTs. Specially, the fractional abundances of most isoGDGTs, except for crenarchaeol', correlate significantly with temperature, whereas only a few brGDGT compounds show a notable relationship with the mean temperature. Among the isoGDGTs, GDGT-0 and crenarchaeol are most strongly correlated with the mean temperature, with Pearson correlation coefficients (*r*) of 0.89 and –0.9, respectively. Among the brGDGTs, IIa' and Ia show the highest positive correlations, with *r* values of 0.73 and 0.6, respectively. (b) Correlation coefficients (*r*) between the GDGT-based proxies and corresponding interval mean temperature. Among the GDGT-based proxies, the RI ($r$ = –0.9) demonstrated the greatest sensitivity to temperature, outperforming $TEX_{86}$ ($r$ = 0.51). Overall, isoGDGT-based proxies are more sensitive to the mean temperature than brGDGT-based proxies, with higher correlations observed at shallow depths compared to deeper waters. Significant correlations ($P \leq 0.05$) are indicated with asterisks.

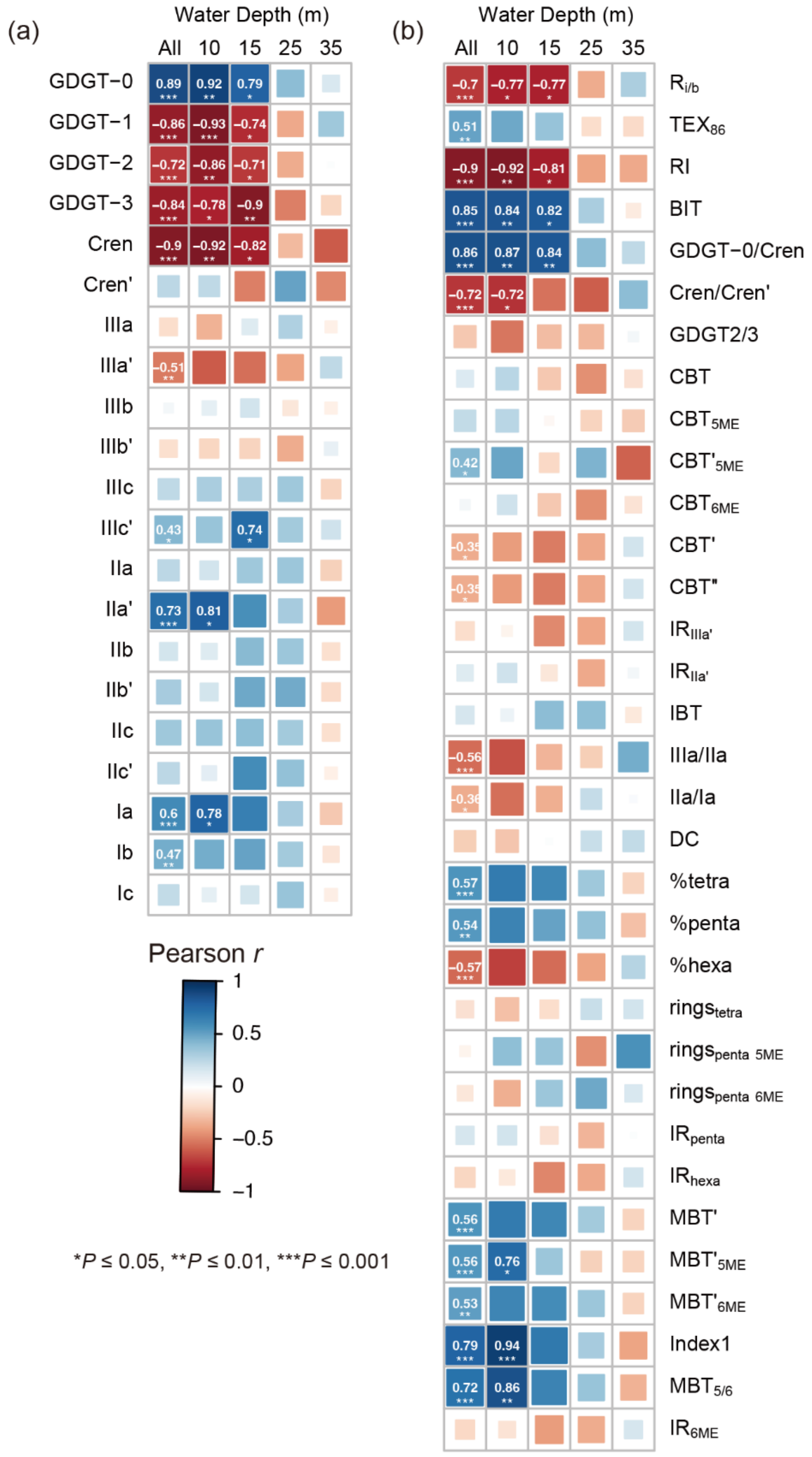
(a)
Water Depth (m)
All 10 15 25 35
GDGT−0
GDGT−1
GDGT−2
GDGT−3
Cren
Cren'
IIIa
IIIa'
IIIb
IIIb'
IIIc
IIIc'
IIa
IIa'
IIb
IIb'
IIc
IIc'
Ia
Ib
Ic
Pearson r
1
0.5
0
−0.5
−1
*P ≤ 0.05, **P ≤ 0.01, ***P ≤ 0.001
(b)
Water Depth (m)
All 10 15 25 35
Ri/b
TEX86
RI
BIT
GDGT−0/Cren
Cren/Cren'
GDGT2/3
CBT
CBT5ME
CBT'5ME
CBT6ME
CBT'
CBT"
IRIIIa'
IRIIa'
IBT
IIIa/IIa
IIa/Ia
DC
%tetra
%penta
%hexa
ringstetra
ringspenta 5ME
ringspenta 6ME
IRpenta
IRhexa
MBT'
MBT'5ME
MBT'6ME
Index1
MBT5/6
IR6ME

**Fig. S5.** Correlation analysis between the daily mean temperature (DMT) and selected GDGT components. Scatter plots show the relationships between DMT over the 60 days preceding trap recovery, and (a) GDGT-0, (b) crenarchaeol, and (c) the RI. The trap recovery date is defined as day 0, with negative days representing the period prior to trap recovery (day –1 to day –60). Day 0 corresponds to the daily mean temperature measured across all trap deployment depths on the trap recovery date.

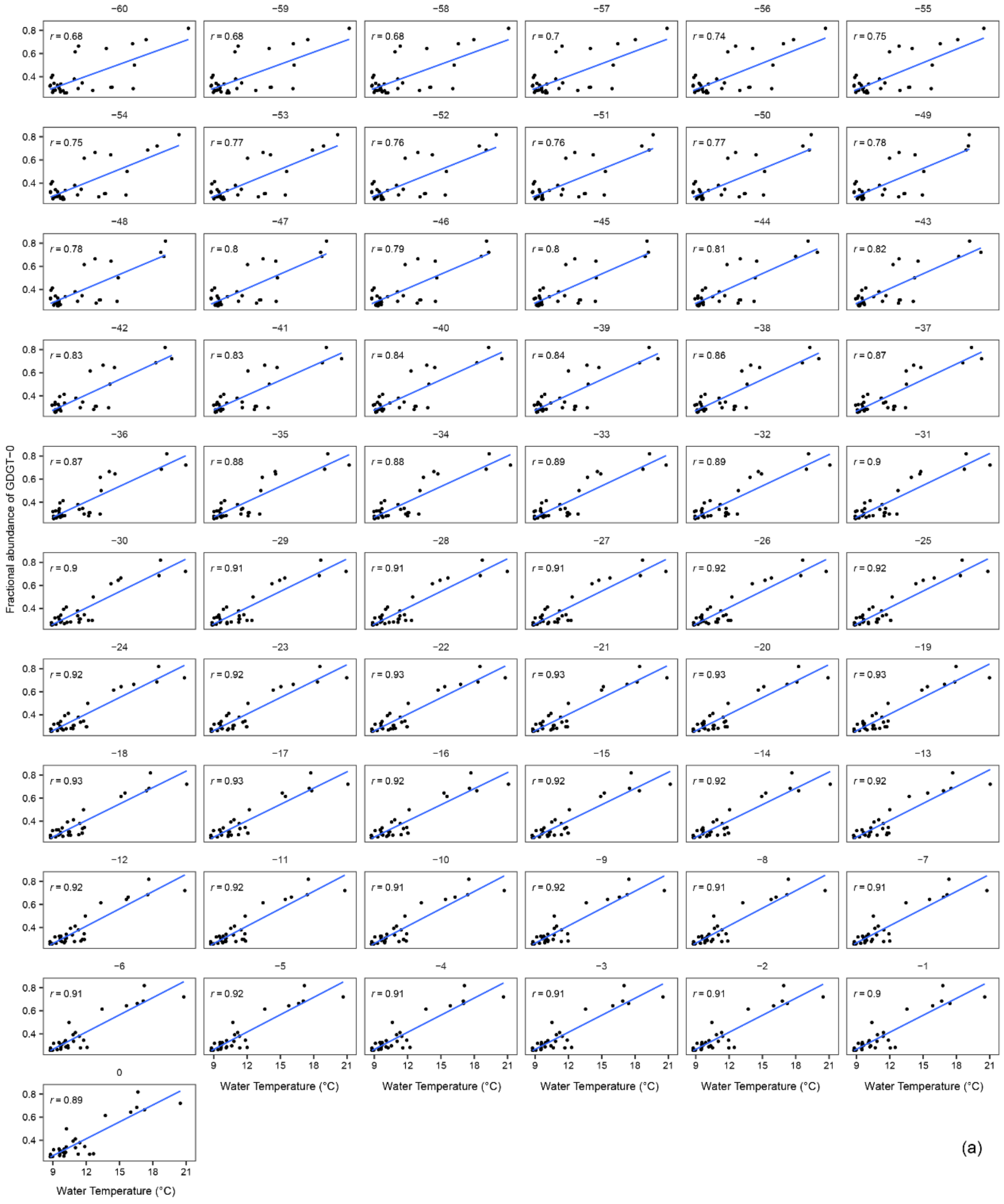

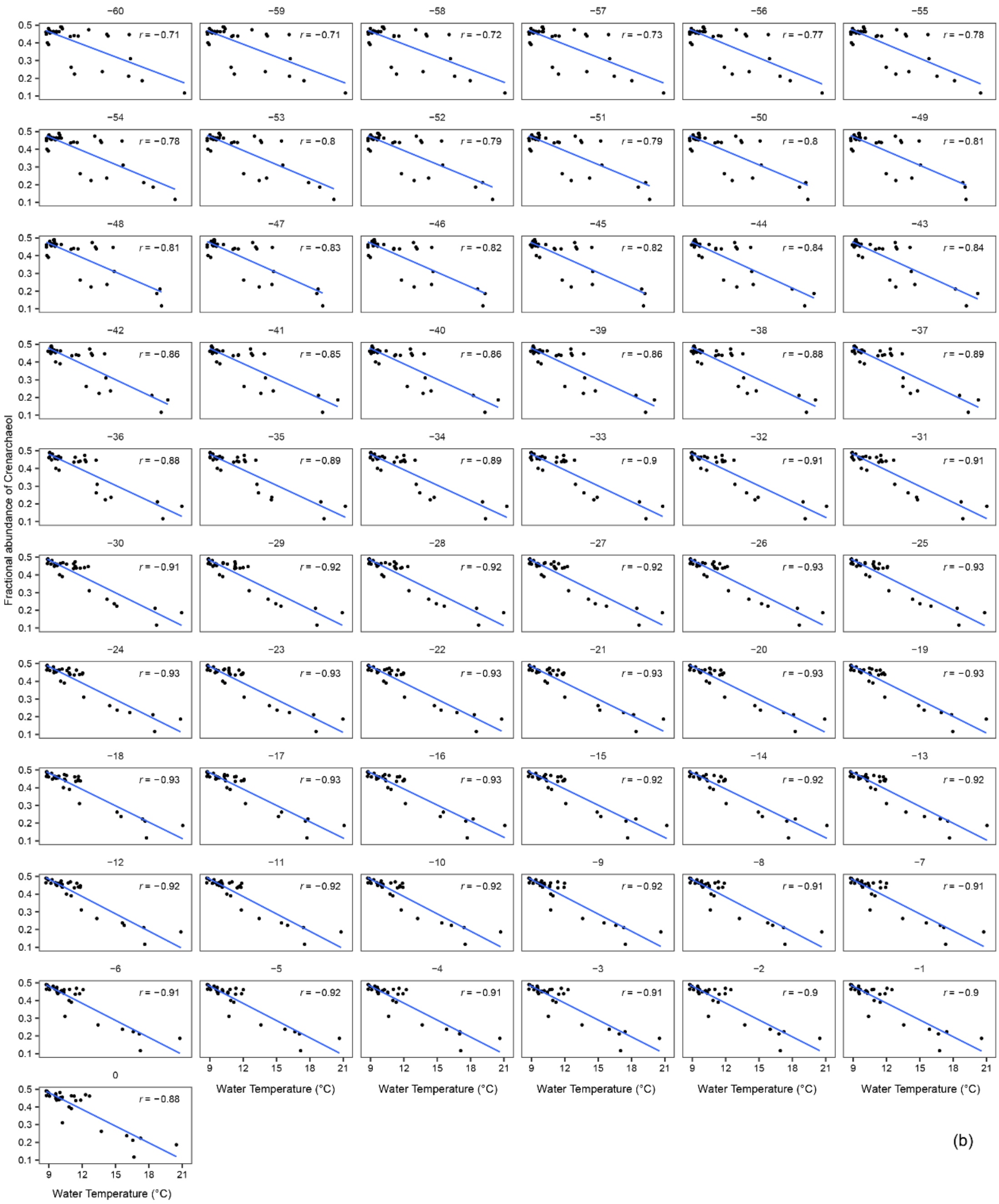
Fractional abundance of Crenarchaeol
Water Temperature (°C)
−60 r = −0.71
−59 r = −0.71
−58 r = −0.72
−57 r = −0.73
−56 r = −0.77
−55 r = −0.78
−54 r = −0.78
−53 r = −0.8
−52 r = −0.79
−51 r = −0.79
−50 r = −0.8
−49 r = −0.81
−48 r = −0.81
−47 r = −0.83
−46 r = −0.82
−45 r = −0.82
−44 r = −0.84
−43 r = −0.84
−42 r = −0.86
−41 r = −0.85
−40 r = −0.86
−39 r = −0.86
−38 r = −0.88
−37 r = −0.89
−36 r = −0.88
−35 r = −0.89
−34 r = −0.89
−33 r = −0.9
−32 r = −0.91
−31 r = −0.91
−30 r = −0.91
−29 r = −0.92
−28 r = −0.92
−27 r = −0.92
−26 r = −0.93
−25 r = −0.93
−24 r = −0.93
−23 r = −0.93
−22 r = −0.93
−21 r = −0.93
−20 r = −0.93
−19 r = −0.93
−18 r = −0.93
−17 r = −0.93
−16 r = −0.93
−15 r = −0.92
−14 r = −0.92
−13 r = −0.92
−12 r = −0.92
−11 r = −0.92
−10 r = −0.92
−9 r = −0.92
−8 r = −0.91
−7 r = −0.91
−6 r = −0.91
−5 r = −0.92
−4 r = −0.91
−3 r = −0.91
−2 r = −0.9
−1 r = −0.9
0 r = −0.88
(b)

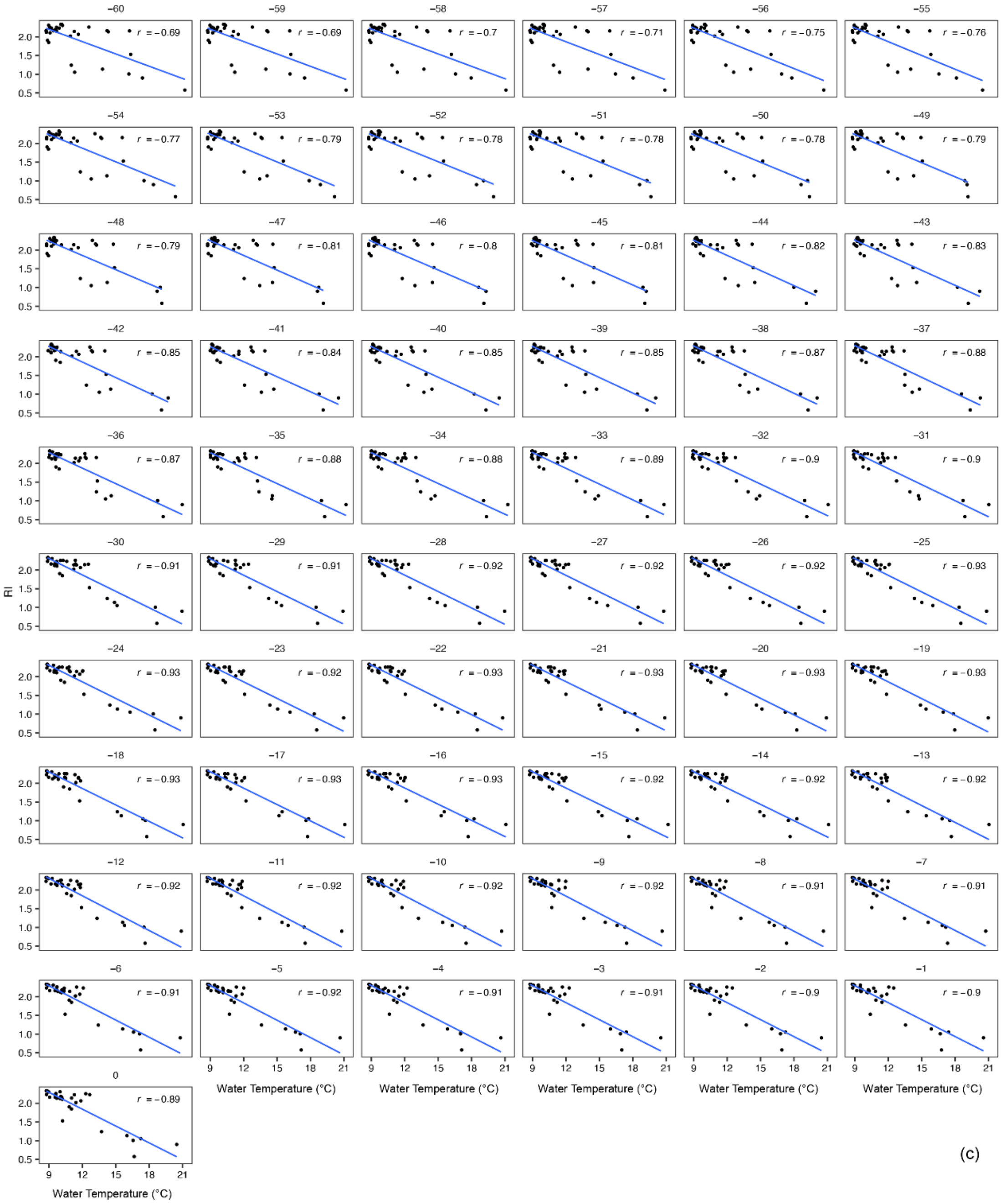

RI
Water Temperature (°C)
2.0
1.5
1.0
0.5
9
12
15
18
21
−60
r = −0.69
−59
r = −0.69
−58
r = −0.7
−57
r = −0.71
−56
r = −0.75
−55
r = −0.76
−54
r = −0.77
−53
r = −0.79
−52
r = −0.78
−51
r = −0.78
−50
r = −0.78
−49
r = −0.79
−48
r = −0.79
−47
r = −0.81
−46
r = −0.8
−45
r = −0.81
−44
r = −0.82
−43
r = −0.83
−42
r = −0.85
−41
r = −0.84
−40
r = −0.85
−39
r = −0.85
−38
r = −0.87
−37
r = −0.88
−36
r = −0.87
−35
r = −0.88
−34
r = −0.88
−33
r = −0.89
−32
r = −0.9
−31
r = −0.9
−30
r = −0.91
−29
r = −0.91
−28
r = −0.92
−27
r = −0.92
−26
r = −0.92
−25
r = −0.93
−24
r = −0.93
−23
r = −0.92
−22
r = −0.93
−21
r = −0.93
−20
r = −0.93
−19
r = −0.93
−18
r = −0.93
−17
r = −0.93
−16
r = −0.93
−15
r = −0.92
−14
r = −0.92
−13
r = −0.92
−12
r = −0.92
−11
r = −0.92
−10
r = −0.92
−9
r = −0.92
−8
r = −0.91
−7
r = −0.91
−6
r = −0.91
−5
r = −0.92
−4
r = −0.91
−3
r = −0.91
−2
r = −0.9
−1
r = −0.9
0
r = −0.89
(c)

**Fig. S6.** Pearson correlation analysis between DMT and GDGT distributions in settling particles. Panels (a) and (c) show the variability in Pearson *r* values between DMT over the 60 days preceding trap recovery and the fractional abundance of GDGT-0 and RI, respectively. Correlations with |*r*| < 0.8 are shown as grey dots. Panels (b) and (d) display scatter plots illustrating the relationship between DMT on day –20 ($DMT_{-20}$) and the fractional abundance of GDGT-0 and RI. $DMT_{-20}$ yields the maximum absolute correlation (|*r*|), highlighted with black circles. Panels (e) and (f) show circos plots visualizing the correlation between DMT and individual GDGTs and GDGT-based proxies. The upper quarter of each circos plot highlights the four variables with the strongest correlations (|*r*| > 0.8). The top arc comprises 31 data points representing Pearson *r* for DMT values averaged over consecutive two-day intervals within the 60 days before trap recovery. Proxy definitions are provided in Tables 1 and S1.

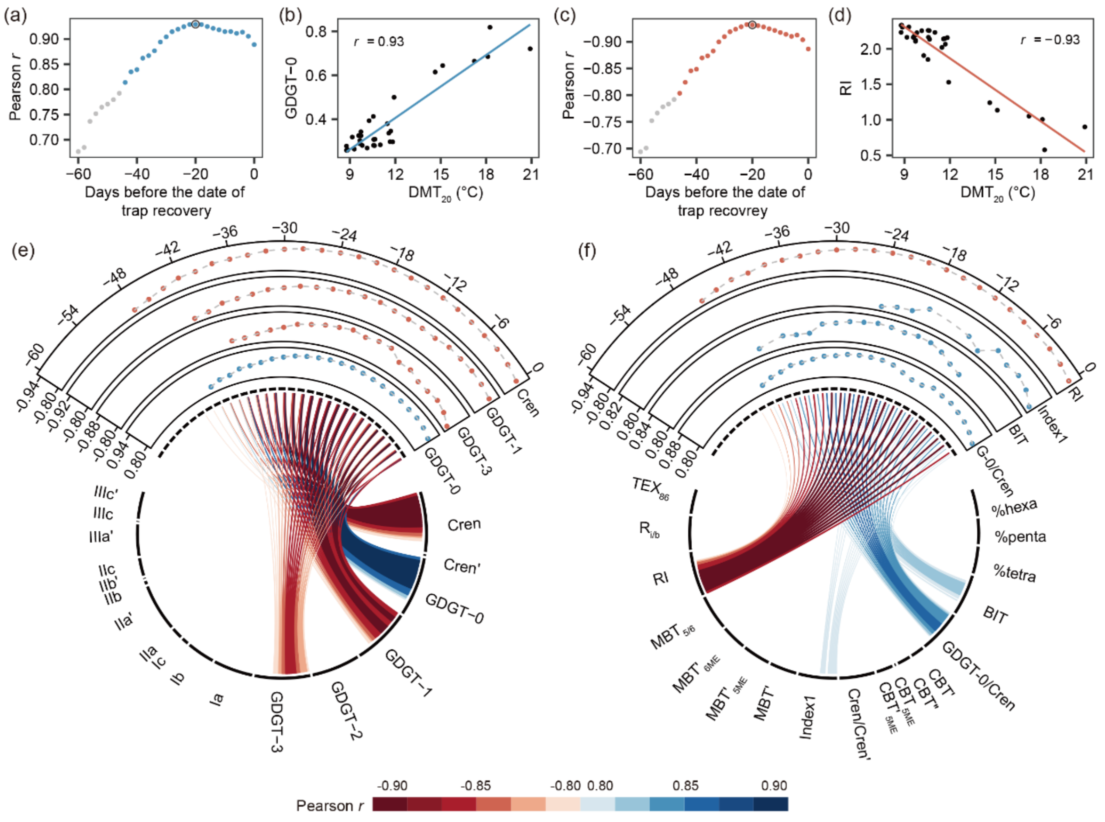

**Fig. S7.** Pearson correlation coefficients between DMT and (a) the fractional abundance of GDGTs and (b) GDGT-based proxies. DMT was evaluated over the 60 days prior to trap recovery. Day 0 denotes the trap recovery date, while day –60 refers to 60 days before trap recovery.

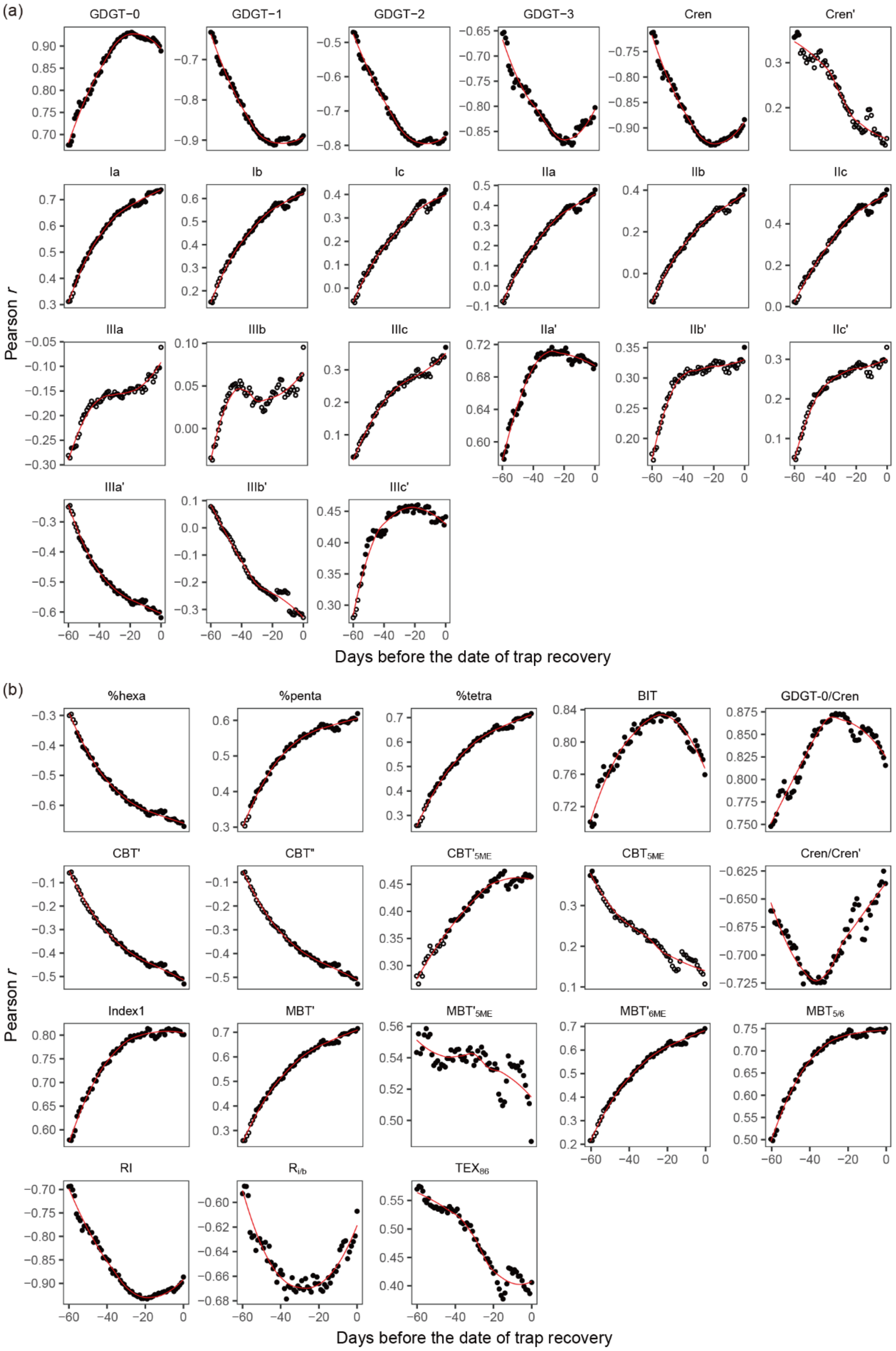
(a)
GDGT−0
GDGT−1
GDGT−2
GDGT−3
Cren
Cren'
Ia
Ib
Ic
IIa
IIb
IIc
IIIa
IIIb
IIIc
IIa'
IIb'
IIc'
IIIa'
IIIb'
IIIc'
Pearson r
Days before the date of trap recovery
(b)
%hexa
%penta
%tetra
BIT
GDGT-0/Cren
CBT'
CBT''
CBT'5ME
CBT5ME
Cren/Cren'
Index1
MBT'
MBT'5ME
MBT'6ME
MBT5/6
RI
RI/b
TEX86
Pearson r
Days before the date of trap recovery

**Fig. S8.** Relationships between the incubation temperature and GDGT-based proxies in Thaumarchaeota cultures. (a) Scatter plot showing the relationship between incubation temperature and $TEX_{86}$ for Thaumarchaeota Group I.1a (blue) and I.1b (pink), with an overall $R^2$ of 0.56. (b) Relationship between incubation temperature and the RI, with an $R^2$ of 0.86. When only Group I.1a is considered, $R^2$ values decrease to 0.24 for $TEX_{86}$ and 0.73 for RI. Data sources compiled from Pitcher et al. (2010); (2011); Sinninghe Damsté et al. (2012); Elling et al. (2015); Qin et al. (2015); Elling et al. (2017). Blue regression lines indicate relationships based on Group I.1a cultures only.

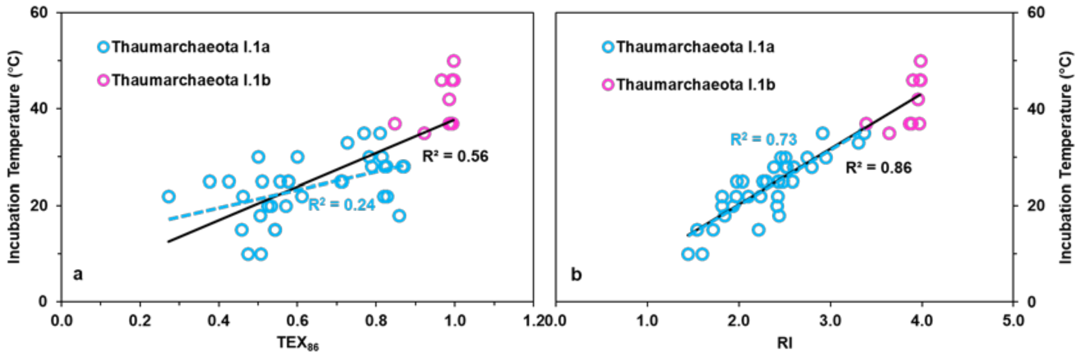

**Fig. S9.** Relationship between RI and $TEX_{86}$ across various sedimentary archives and particulate samples. The dataset includes settling particles from this study; global marine surface sediments (Kim et al., 2010); lake surface sediments (Günther et al., 2014; Hu et al., 2016; Wang et al., 2016b; Wang et al., 2020); lake suspended particulate matter (SPM) (Hu et al., 2016) and settling particles (Hu et al., 2016); soils (Wang et al., 2012; Yang et al., 2012; Liu et al., 2013; Hu et al., 2016; Li et al., 2016; Wang et al., 2016b; Yao et al., 2019); and pure cultures of Thaumarchaeota Groups I.1a and I.1b (Pitcher et al., 2010; Pitcher et al., 2011; Sinninghe Damsté et al., 2012; Elling et al., 2015; Qin et al., 2015; Elling et al., 2017). Strong linear correlations between RI and $TEX_{86}$ were only observed in global marine surface sediments ($R^2$ = 0.87, navy) and Thaumarchaeota cultures ($R^2$ = 0.60, purple), suggesting environmental or source-specific constraints on proxy coherence.

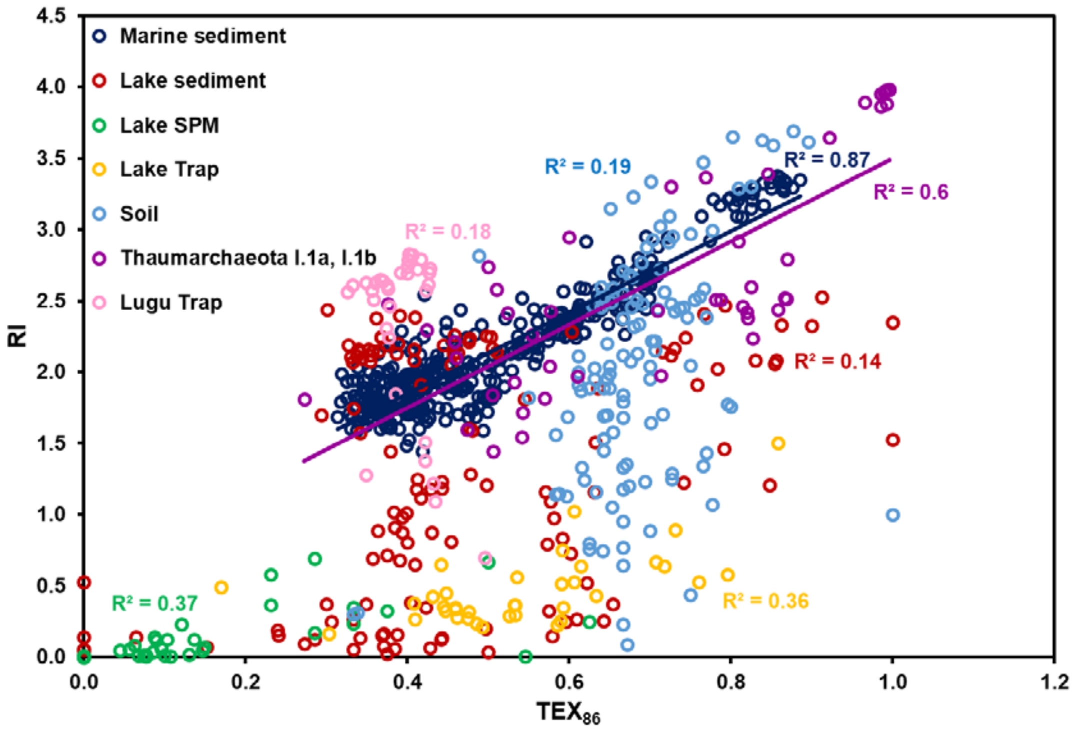

**Fig. S10.** Ternary diagram illustrating the fractional abundances of summed tetramethylated, pentamethylated, and hexamethylated brGDGTs. The dataset includes settling particles from this study and lake surface sediments from a global compilation of lakes reported by Martínez-Sosa et al. (2021).

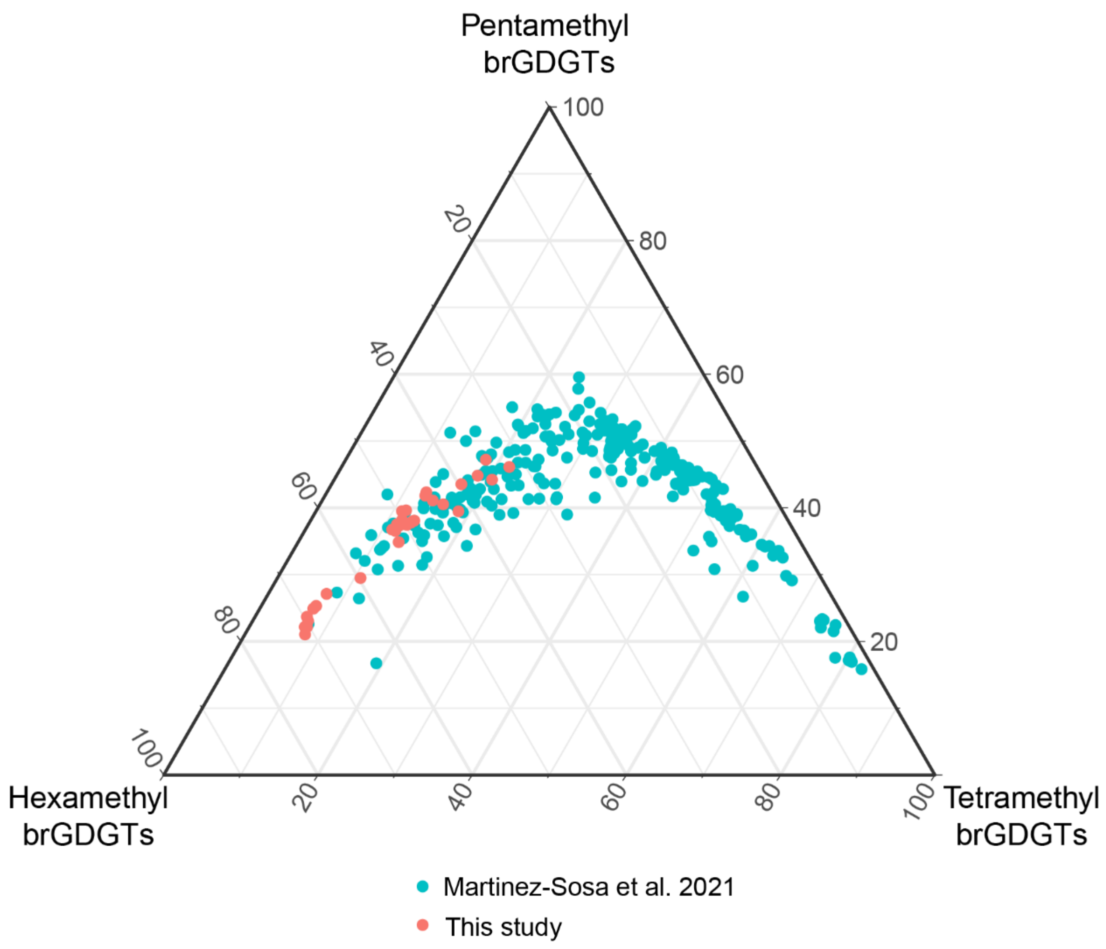

**Fig. S11.** Variability in Pearson correlation coefficients between RI and multiple-day mean temperatures (MMT) across different window sizes. Pearson correlations coefficients were calculated between RI and MMT using sliding windows ranging from 1 to 30 days. An increase in window size leads to a corresponding reduction in the number of available overlapping windows. For each window size, the correlation was computed by incrementally shifting the window by one day throughout the sampling period. The first panel ($MMT_1$) corresponds to daily mean temperature (DMT), replicating the correlation patterns of RI shown in Fig. S7b. In each panel, the highest absolute Pearson correlation coefficient (|$r$|) is highlighted in red.

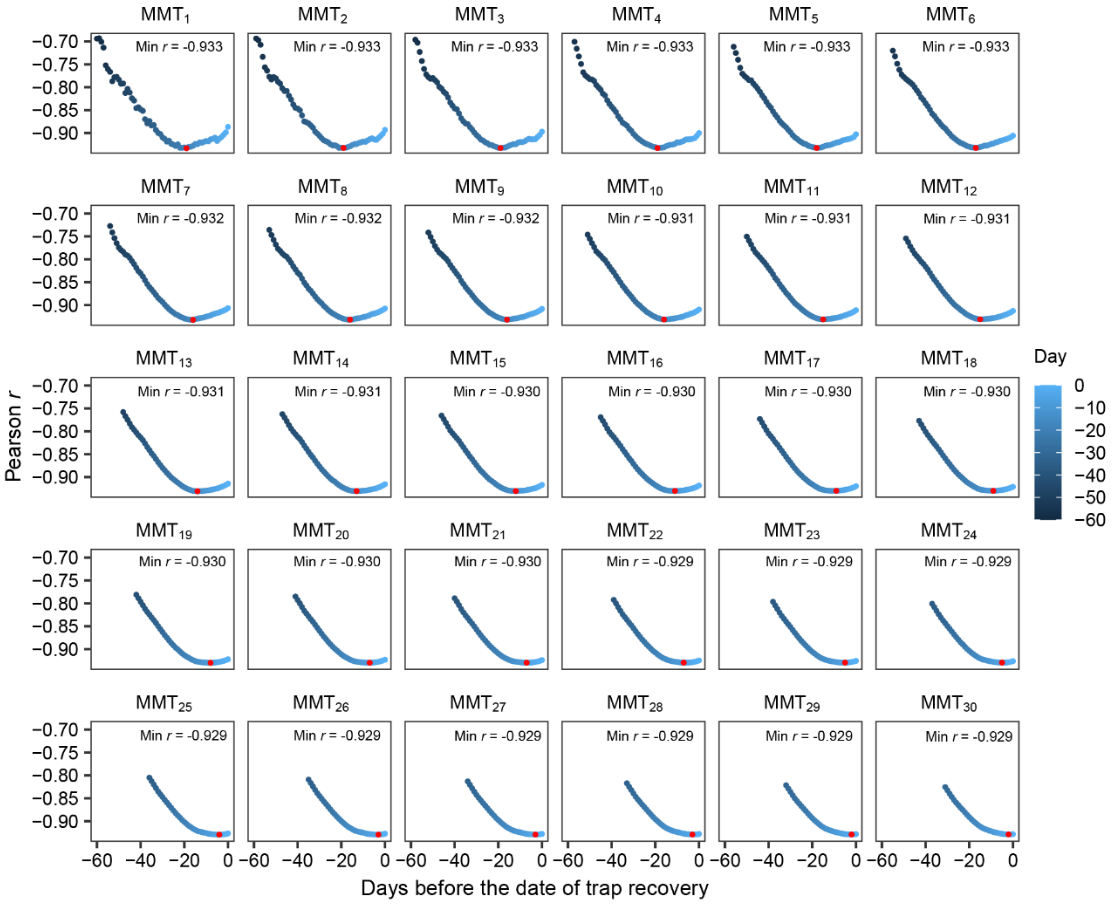

**Fig. S12.** Temporal lags and vertical offsets in the relationship between $MMT_{20}$ and Ia. Ia was calculated from brGDGTs in settling particles collected at four depths (10, 15, 25, and 35 m). Panels (a–d) illustrate the spatiotemporal patterns of Ia–$MMT_{20}$ correlations at each trap depth. Each panel contains three subplots: The upper subpanels show the temporal evolution of the Pearson correlation coefficient (*r*) between Ia and $MMT_{20}$, calculated over 42 overlapping windows prior to trap recovery. The left subpanels display vertical correlation profiles, showing Pearson *r* values between Ia at each trap depth and $MMT_{20}$ calculated at 0.5 m intervals from 0 to 20 m and 2 m intervals from 20 to 40 m. The central subpanels summarize these vertical patterns, combining correlations between Ia and $MMT_{20}$ at all depths above the trap depth (blue points), with gray points representing depths below the trap. Fitted red curves indicate loess-smoothed trends in depth-wise correlations.

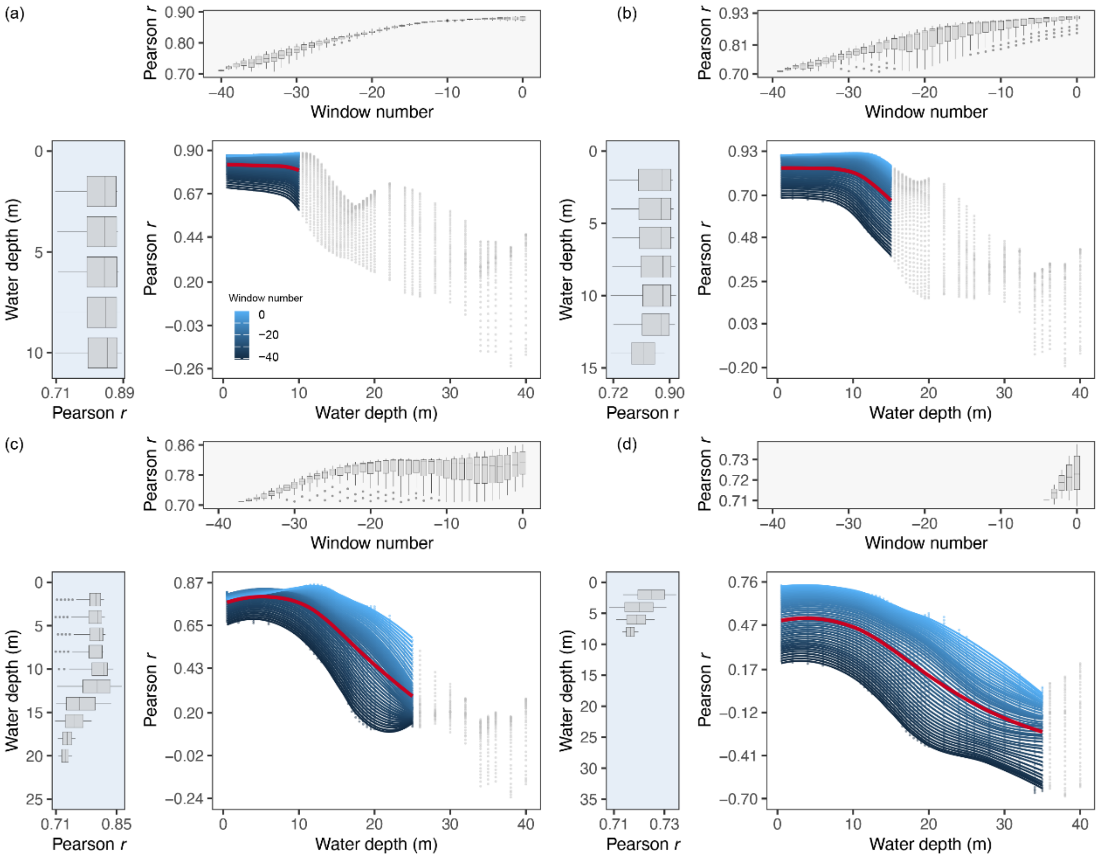

### Supplemental Tables

**Tab. S1.** Equations for GDGT-based proxies used in this study.

| **NO.** | **Equation** | **Reference** |
|---|---|---|
| 1 | $CBT_{5ME}=^{-10}log((Ib+IIb)/(Ia+IIa))$ | De Jonge et al. (2014) |
| 2 | $CBT'_{5ME}=-log(IIb/IIa)$ | Wang et al. (2016a) |
| 3 | $CBT_{6ME}=-log((Ib+IIb')/(Ia+IIa'))$ | Yang et al. (2015) |
| 4 | CBT'=log[(Ic+IIa'+IIb'+IIc'+IIIa'+IIIb'+IIIc')/(Ia+IIa+IIIa)] | De Jonge et al. (2014) |
| 5 | CBT''=log[(Ib+Ic+IIb+IIc+IIIb+IIIc+IIa'+IIb'+IIc'+IIIa'+IIIb'+IIIc')/(Ia+IIa+IIIa)] | Ding et al. (2015) |
| 6 | $IR_{IIIa}$=IIIa'/(IIIa'+IIIa) | De Jonge et al. (2014) |
| 7 | $IR_{IIa}$=IIa'/(IIa'+IIa) | De Jonge et al. (2014) |
| 8 | IBT=-log((IIIa'+IIa')/(IIIa+IIa)) | Ding et al. (2015) |
| 9 | DC=(Ib+IIb)/(Ia+IIa+Ib+IIb) | Sinninghe Damsté et al. (2009) |
| 10 | %tetra= 100*(Ia+Ib+Ic)/ΣbrGDGTs | Sinninghe Damsté (2016) |
| 11 | %penta=100*(IIa+IIb+IIc+IIa'+IIb'+IIc') /ΣbrGDGTs | Sinninghe Damsté (2016) |
| 12 | %hexa=100*(IIIa+IIIb+IIIc+IIIa'+IIIb'+IIIc') /ΣbrGDGTs | Sinninghe Damsté (2016) |
| 13 | $rings_{tetra}$=(Ib+2*Ic)/(Ia+Ib+Ic) | Sinninghe Damsté (2016) |
| 14 | $rings_{penta\ 5ME}$=(IIb+2*IIc)/(IIa+IIb+IIc) | Sinninghe Damsté (2016) |
| 15 | $rings_{penta\ 6ME}$=(IIb'+2*IIc')/(IIa'+IIb'+IIc') | Sinninghe Damsté (2016) |
| 16 | $IR_{penta}$=(IIa'+IIb'+IIc')/all penta brGDGTs | Sinninghe Damsté (2016) |
| 17 | $IR_{hexa}$=(IIIa'+IIIb'+IIIc')/all hexa brGDGTs | Sinninghe Damsté (2016) |
| 18 | MBT'=(Ia+Ib+Ic)/(Ia+Ib+Ic+IIa+IIb+IIc+IIIa+IIa'+IIb'+IIc'+IIIa') | Peterse et al. (2012) |
| 19 | $MBT_{5/6}$=(Ia+Ib+Ic+IIa')/(Ia+Ib+Ic+IIa+IIb+IIc+IIIa+IIIa') | Ding et al. (2015) |
| 20 | MI=(GDGT-1+GDGT-2+GDGT-3)/(GDGT-1+GDGT-2+GDGT-3+Cren+Cren') | Zhang et al. (2011) |

**Tab. S2.** Fluxes and GDGT-based proxies in settling particles collected at four water depths in Lake Lugu from October 2012 and April 2014.

| Sample Code | Depth | Deployment date | Recovery date | Mean temp. | Min. temp. | Max. temp. | Mass flux | ΣisoGDGTs | ΣbrGDGTs | GDGT-0 | GDGT-1 | GDGT-2 | GDGT-3 | Cren | Cren' |
|---|---|---|---|---|---|---|---|---|---|---|---|---|---|---|---|
| Name-depth-recovery year/m | m | yyyy-mm-dd | | temperature of each collecting period (°C) | | | g/m$^2$/day | μg/m$^2$/day | | ng/m$^2$/day | | | | | |
| TP2-10-1212 | 10 | 2012-10-25 | 2012-12-29 | 13.3 | 10.2 | 17.0 | 0.04 | 0.02 | 0.07 | 10 | 2 | 1 | 0 | 6 | 0 |
| TP2-10-1302 | 10 | 2012-12-29 | 2013-02-27 | 9.2 | 8.8 | 10.4 | 0.74 | 4.25 | 1.88 | 1107 | 661 | 378 | 45 | 2037 | 20 |
| TP2-10-1304 | 10 | 2013-02-27 | 2013-04-18 | 10.4 | 8.9 | 11.4 | 0.22 | 0.19 | 0.22 | 78 | 23 | 12 | 2 | 74 | 1 |
| TP2-10-1306 | 10 | 2013-04-18 | 2013-06-19 | 14.9 | 11.0 | 18.4 | 0.06 | 0.02 | 0.06 | 12 | 1 | 1 | 0 | 4 | 0 |
| TP2-10-1308 | 10 | 2013-06-19 | 2013-08-22 | 20.0 | 17.2 | 21.3 | 0.16 | 0.04 | 0.11 | 25 | 2 | 1 | 0 | 7 | 0 |
| TP2-10-1310 | 10 | 2013-08-22 | 2013-10-27 | 18.9 | 16.7 | 21.2 | 0.25 | 0.04 | 0.46 | 35 | 1 | 1 | 0 | 5 | 0 |
| TP2-10-1401 | 10 | 2013-10-27 | 2014-01-17 | 12.7 | 9.6 | 16.9 | 0.69 | 1.30 | 1.26 | 402 | 190 | 120 | 13 | 571 | 6 |
| TP2-10-1404 | 10 | 2014-01-17 | 2014-04-28 | 10.1 | 9.1 | 12.7 | 0.66 | 3.26 | 1.30 | 924 | 478 | 302 | 36 | 1504 | 18 |
| TP3-15-1212 | 15 | 2012-10-25 | 2012-12-29 | 13.2 | 10.1 | 16.9 | 0.24 | 2.28 | 1.19 | 678 | 366 | 189 | 23 | 1021 | 8 |
| TP3-15-1302 | 15 | 2012-12-29 | 2013-02-27 | 9.1 | 8.7 | 10.3 | 0.78 | 4.67 | 1.98 | 1208 | 700 | 406 | 49 | 2281 | 23 |
| TP3-15-1304 | 15 | 2013-02-27 | 2013-04-18 | 10.0 | 8.8 | 11.0 | 0.45 | 0.46 | 0.36 | 182 | 60 | 29 | 5 | 186 | 2 |
| TP3-15-1306 | 15 | 2013-04-18 | 2013-06-19 | 13.1 | 10.8 | 15.5 | 0.07 | 0.03 | 0.07 | 19 | 2 | 1 | 0 | 8 | 0 |
| TP3-15-1308 | 15 | 2013-06-19 | 2013-08-22 | 14.8 | 13.7 | 16.0 | 0.10 | 0.03 | 0.07 | 19 | 2 | 1 | 0 | 7 | 0 |
| TP3-15-1310 | 15 | 2013-08-22 | 2013-10-27 | 17.8 | 15.9 | 19.3 | 0.23 | 0.06 | 0.20 | 38 | 3 | 2 | 0 | 12 | 0 |
| TP3-15-1401 | 15 | 2013-10-27 | 2014-01-17 | 12.6 | 9.5 | 16.8 | 0.66 | 1.17 | 1.20 | 359 | 163 | 102 | 12 | 522 | 6 |
| TP3-15-1404 | 15 | 2014-01-17 | 2014-04-28 | 9.8 | 9.0 | 12.3 | 1.06 | 5.87 | 2.05 | 1644 | 843 | 538 | 65 | 2755 | 29 |
| TP3-25-1212 | 25 | 2012-10-25 | 2012-12-29 | 11.4 | 9.9 | 12.2 | 0.11 | 1.02 | 0.53 | 304 | 167 | 86 | 11 | 451 | 4 |
| TP3-25-1302 | 25 | 2012-12-29 | 2013-02-27 | 9.1 | 8.7 | 10.3 | 0.68 | 3.59 | 1.28 | 998 | 565 | 315 | 38 | 1667 | 11 |
| TP3-25-1304 | 25 | 2013-02-27 | 2013-04-18 | 9.4 | 8.8 | 10.1 | 0.38 | 1.25 | 0.57 | 405 | 182 | 83 | 13 | 560 | 5 |

| | | | | | | | | | | | | | | | |
|---|---|---|---|---|---|---|---|---|---|---|---|---|---|---|---|
| TP3-25-1306 | 25 | 2013-04-18 | 2013-06-19 | 10.8 | 9.7 | 11.7 | 0.16 | 0.66 | 0.34 | 221 | 88 | 34 | 7 | 303 | 3 |
| TP3-25-1308 | 25 | 2013-06-19 | 2013-08-22 | 11.2 | 10.8 | 11.5 | 0.20 | 0.34 | 0.28 | 131 | 40 | 19 | 4 | 150 | 2 |
| TP3-25-1310 | 25 | 2013-08-22 | 2013-10-27 | 11.7 | 11.4 | 12.0 | 0.36 | 0.96 | 0.44 | 333 | 133 | 61 | 10 | 422 | 4 |
| TP3-25-1401 | 25 | 2013-10-27 | 2014-01-17 | 11.6 | 9.5 | 13.1 | 0.63 | 3.53 | 1.23 | 991 | 519 | 292 | 37 | 1673 | 17 |
| TP3-25-1404 | 25 | 2014-01-17 | 2014-04-28 | 9.5 | 9.0 | 11.3 | 0.79 | 3.18 | 1.28 | 892 | 493 | 278 | 35 | 1465 | 16 |
| TP2-35-1212 | 35 | 2012-10-25 | 2012-12-29 | 9.7 | 9.5 | 10.2 | 0.18 | 1.61 | 0.90 | 465 | 262 | 132 | 16 | 730 | 5 |
| TP2-35-1302 | 35 | 2012-12-29 | 2013-02-27 | 9.1 | 8.7 | 10.2 | 1.00 | 7.70 | 3.12 | 1990 | 1156 | 665 | 79 | 3774 | 34 |
| TP2-35-1304 | 35 | 2013-02-27 | 2013-04-18 | 9.0 | 8.8 | 9.2 | 0.76 | 2.86 | 1.21 | 912 | 403 | 187 | 29 | 1319 | 11 |
| TP2-35-1306 | 35 | 2013-04-18 | 2013-06-19 | 9.6 | 9.1 | 10.3 | 0.18 | 0.82 | 0.45 | 281 | 112 | 43 | 8 | 373 | 2 |
| TP2-35-1308 | 35 | 2013-06-19 | 2013-08-22 | 9.5 | 9.4 | 10.2 | 0.15 | 0.54 | 0.25 | 176 | 74 | 31 | 6 | 251 | 2 |
| TP2-35-1310 | 35 | 2013-08-22 | 2013-10-27 | 9.6 | 9.5 | 9.7 | 0.31 | 2.75 | 0.68 | 833 | 410 | 198 | 28 | 1269 | 11 |
| TP2-35-1401 | 35 | 2013-10-27 | 2014-01-17 | 9.8 | 9.5 | 10.4 | 0.98 | 21.73 | 4.10 | 5839 | 3401 | 2079 | 223 | 10098 | 91 |
| TP2-35-1404 | 35 | 2014-01-17 | 2014-04-28 | 9.2 | 8.9 | 10.0 | 1.20 | 8.10 | 2.39 | 2137 | 1202 | 731 | 88 | 3896 | 41 |

| Sample Code | Depth | IIIa | IIIa' | IIIb | IIIb' | IIIc | IIIc' | IIa | IIa' | IIb | IIb' | IIc | IIc' | Ia | Ib | Ic |
|---|---|---|---|---|---|---|---|---|---|---|---|---|---|---|---|---|
| | m | | | | | | | | ng/m$^2$/day | | | | | | | |
| TP2-10-1212 | 10 | 3 | 26 | 0 | 1 | 0 | 0 | 3 | 18 | 2 | 4 | 0 | 0 | 6 | 2 | 1 |
| TP2-10-1302 | 10 | 40 | 1267 | 4 | 11 | 2 | 1 | 46 | 290 | 31 | 43 | 3 | 4 | 107 | 21 | 7 |
| TP2-10-1304 | 10 | 9 | 94 | 1 | 1 | 0 | 0 | 14 | 47 | 10 | 11 | 1 | 1 | 21 | 7 | 2 |
| TP2-10-1306 | 10 | 2 | 19 | 0 | 0 | 0 | 0 | 4 | 14 | 3 | 3 | 0 | 0 | 6 | 3 | 1 |
| TP2-10-1308 | 10 | 4 | 34 | 1 | 0 | 0 | 0 | 8 | 30 | 5 | 6 | 1 | 1 | 17 | 5 | 1 |
| TP2-10-1310 | 10 | 8 | 211 | 1 | 2 | 1 | 0 | 14 | 138 | 9 | 18 | 2 | 1 | 44 | 8 | 1 |
| TP2-10-1401 | 10 | 64 | 537 | 6 | 6 | 4 | 1 | 59 | 311 | 40 | 73 | 6 | 7 | 101 | 32 | 7 |
| TP2-10-1404 | 10 | 69 | 574 | 6 | 7 | 4 | 2 | 72 | 289 | 50 | 63 | 6 | 6 | 107 | 34 | 8 |

| | | | | | | | | | | | | | | | | |
|---|---|---|---|---|---|---|---|---|---|---|---|---|---|---|---|---|
| TP3-15-1212 | 15 | 41 | 748 | 8 | 9 | 2 | 1 | 37 | 186 | 28 | 39 | 3 | 3 | 62 | 16 | 4 |
| TP3-15-1302 | 15 | 42 | 1331 | 4 | 12 | 3 | 1 | 45 | 311 | 31 | 45 | 3 | 4 | 120 | 24 | 5 |
| TP3-15-1304 | 15 | 17 | 140 | 2 | 2 | 1 | 0 | 26 | 84 | 17 | 19 | 2 | 1 | 36 | 13 | 3 |
| TP3-15-1306 | 15 | 3 | 21 | 0 | 0 | 0 | 0 | 6 | 20 | 4 | 5 | 1 | 0 | 10 | 3 | 1 |
| TP3-15-1308 | 15 | 3 | 17 | 0 | 0 | 0 | 0 | 5 | 17 | 4 | 4 | 1 | 0 | 11 | 3 | 1 |
| TP3-15-1310 | 15 | 10 | 60 | 1 | 1 | 0 | 1 | 13 | 52 | 9 | 11 | 1 | 1 | 27 | 8 | 1 |
| TP3-15-1401 | 15 | 73 | 445 | 8 | 5 | 5 | 2 | 70 | 293 | 49 | 82 | 6 | 7 | 106 | 40 | 9 |
| TP3-15-1404 | 15 | 129 | 878 | 13 | 12 | 6 | 3 | 128 | 424 | 86 | 107 | 10 | 8 | 166 | 62 | 14 |
| TP3-25-1212 | 25 | 19 | 335 | 2 | 4 | 1 | 1 | 17 | 84 | 13 | 18 | 1 | 2 | 28 | 8 | 2 |
| TP3-25-1302 | 25 | 31 | 861 | 6 | 8 | 2 | 1 | 25 | 192 | 23 | 24 | 2 | 3 | 78 | 17 | 4 |
| TP3-25-1304 | 25 | 27 | 244 | 3 | 3 | 2 | 1 | 33 | 125 | 24 | 29 | 3 | 3 | 55 | 16 | 4 |
| TP3-25-1306 | 25 | 20 | 143 | 2 | 1 | 1 | 1 | 22 | 69 | 16 | 16 | 2 | 2 | 31 | 10 | 2 |
| TP3-25-1308 | 25 | 13 | 100 | 1 | 1 | 1 | 1 | 19 | 57 | 14 | 17 | 2 | 1 | 37 | 12 | 2 |
| TP3-25-1310 | 25 | 24 | 161 | 2 | 2 | 1 | 1 | 28 | 100 | 20 | 25 | 2 | 2 | 53 | 14 | 3 |
| TP3-25-1401 | 25 | 85 | 507 | 9 | 7 | 4 | 2 | 73 | 249 | 54 | 79 | 6 | 7 | 99 | 37 | 9 |
| TP3-25-1404 | 25 | 88 | 554 | 8 | 8 | 4 | 2 | 81 | 244 | 57 | 72 | 7 | 7 | 104 | 38 | 9 |
| TP2-35-1212 | 35 | 27 | 585 | 3 | 7 | 2 | 1 | 25 | 137 | 19 | 27 | 2 | 2 | 45 | 12 | 3 |
| TP2-35-1302 | 35 | 76 | 2067 | 7 | 20 | 4 | 2 | 82 | 486 | 60 | 78 | 7 | 7 | 176 | 38 | 10 |
| TP2-35-1304 | 35 | 61 | 528 | 6 | 6 | 4 | 1 | 71 | 263 | 51 | 60 | 6 | 5 | 112 | 32 | 7 |
| TP2-35-1306 | 35 | 26 | 184 | 3 | 2 | 1 | 1 | 27 | 94 | 22 | 23 | 2 | 2 | 42 | 14 | 3 |
| TP2-35-1308 | 35 | 13 | 116 | 1 | 1 | 1 | 0 | 15 | 49 | 10 | 12 | 1 | 1 | 24 | 8 | 1 |
| TP2-35-1310 | 35 | 26 | 369 | 2 | 3 | 2 | 1 | 29 | 119 | 22 | 24 | 3 | 3 | 55 | 15 | 3 |
| TP2-35-1401 | 35 | 194 | 2426 | 19 | 25 | 9 | 4 | 154 | 657 | 116 | 156 | 13 | 15 | 224 | 67 | 19 |
| TP2-35-1404 | 35 | 157 | 1046 | 16 | 16 | 7 | 3 | 140 | 481 | 105 | 128 | 13 | 12 | 182 | 70 | 17 |

| Sample Code | Depth | Deployment date | Recovery date | Mean temp. | Min. temp. | Max. temp. | $TEX_{86}$ | BIT | GDGT-0/Cren | RI | Cren/Cren' | CBT | $MBT'_{5ME}$ | $MBT'_{6ME}$ | Index 1 | $IR_{6ME}$ |
|---|---|---|---|---|---|---|---|---|---|---|---|---|---|---|---|---|
| | m | yyyy/mm/dd | | of each collecting period (°C) | | | | | | | | | | | | |
| TP2-10-1212 | 10 | 2012-10-25 | 2012-12-29 | 13.3 | 10.2 | 17.0 | 0.39 | 0.90 | 1.61 | 1.5 | 35 | 0.51 | 0.51 | 0.15 | 0.21 | 0.85 |
| TP2-10-1302 | 10 | 2012-12-29 | 2013-02-27 | 9.2 | 8.8 | 10.4 | 0.40 | 0.46 | 0.54 | 2.3 | 104 | 0.67 | 0.53 | 0.08 | 0.09 | 0.93 |
| TP2-10-1304 | 10 | 2013-02-27 | 2013-04-18 | 10.4 | 8.9 | 11.4 | 0.38 | 0.71 | 1.06 | 1.8 | 114 | 0.48 | 0.46 | 0.16 | 0.15 | 0.82 |
| TP2-10-1306 | 10 | 2013-04-18 | 2013-06-19 | 14.9 | 11.0 | 18.4 | 0.35 | 0.92 | 2.97 | 1.1 | 120 | 0.46 | 0.50 | 0.21 | 0.20 | 0.79 |
| TP2-10-1308 | 10 | 2013-06-19 | 2013-08-22 | 20.0 | 17.2 | 21.3 | 0.43 | 0.93 | 3.86 | 0.9 | 24 | 0.54 | 0.57 | 0.25 | 0.26 | 0.79 |
| TP2-10-1310 | 10 | 2013-08-22 | 2013-10-27 | 18.9 | 16.7 | 21.2 | 0.50 | 0.99 | 7.02 | 0.6 | 26 | 0.74 | 0.62 | 0.13 | 0.23 | 0.91 |
| TP2-10-1401 | 10 | 2013-10-27 | 2014-01-17 | 12.7 | 9.6 | 16.9 | 0.42 | 0.65 | 0.70 | 2.1 | 94 | 0.51 | 0.45 | 0.13 | 0.17 | 0.84 |
| TP2-10-1404 | 10 | 2014-01-17 | 2014-04-28 | 10.1 | 9.1 | 12.7 | 0.43 | 0.42 | 0.61 | 2.2 | 86 | 0.50 | 0.43 | 0.14 | 0.14 | 0.82 |
| TP3-15-1212 | 15 | 2012-10-25 | 2012-12-29 | 13.2 | 10.1 | 16.9 | 0.37 | 0.51 | 0.66 | 2.2 | 135 | 0.53 | 0.43 | 0.08 | 0.09 | 0.89 |
| TP3-15-1302 | 15 | 2012-12-29 | 2013-02-27 | 9.1 | 8.7 | 10.3 | 0.41 | 0.45 | 0.53 | 2.3 | 97 | 0.68 | 0.55 | 0.08 | 0.10 | 0.93 |
| TP3-15-1304 | 15 | 2013-02-27 | 2013-04-18 | 10.0 | 8.8 | 11.0 | 0.37 | 0.62 | 0.98 | 1.9 | 94 | 0.48 | 0.45 | 0.18 | 0.16 | 0.79 |
| TP3-15-1306 | 15 | 2013-04-18 | 2013-06-19 | 13.1 | 10.8 | 15.5 | 0.42 | 0.88 | 2.34 | 1.2 | 63 | 0.48 | 0.50 | 0.23 | 0.23 | 0.76 |
| TP3-15-1308 | 15 | 2013-06-19 | 2013-08-22 | 14.8 | 13.7 | 16.0 | 0.42 | 0.88 | 2.71 | 1.1 | 52 | 0.46 | 0.55 | 0.27 | 0.27 | 0.76 |
| TP3-15-1310 | 15 | 2013-08-22 | 2013-10-27 | 17.8 | 15.9 | 19.3 | 0.43 | 0.93 | 3.24 | 1.0 | 56 | 0.52 | 0.53 | 0.22 | 0.24 | 0.79 |
| TP3-15-1401 | 15 | 2013-10-27 | 2014-01-17 | 12.6 | 9.5 | 16.8 | 0.43 | 0.65 | 0.69 | 2.2 | 82 | 0.44 | 0.44 | 0.16 | 0.17 | 0.80 |
| TP3-15-1404 | 15 | 2014-01-17 | 2014-04-28 | 9.8 | 9.0 | 12.3 | 0.43 | 0.38 | 0.60 | 2.3 | 95 | 0.45 | 0.41 | 0.15 | 0.12 | 0.79 |
| TP3-25-1212 | 25 | 2012-10-25 | 2012-12-29 | 11.4 | 9.9 | 12.2 | 0.38 | 0.52 | 0.67 | 2.1 | 106 | 0.52 | 0.43 | 0.08 | 0.09 | 0.89 |
| TP3-25-1302 | 25 | 2012-12-29 | 2013-02-27 | 9.1 | 8.7 | 10.3 | 0.39 | 0.42 | 0.60 | 2.2 | 158 | 0.67 | 0.55 | 0.08 | 0.10 | 0.92 |
| TP3-25-1304 | 25 | 2013-02-27 | 2013-04-18 | 9.4 | 8.8 | 10.1 | 0.36 | 0.46 | 0.72 | 2.1 | 121 | 0.49 | 0.47 | 0.16 | 0.15 | 0.82 |
| TP3-25-1306 | 25 | 2013-04-18 | 2013-06-19 | 10.8 | 9.7 | 11.7 | 0.33 | 0.48 | 0.73 | 2.1 | 95 | 0.46 | 0.42 | 0.16 | 0.13 | 0.79 |
| TP3-25-1308 | 25 | 2013-06-19 | 2013-08-22 | 11.2 | 10.8 | 11.5 | 0.38 | 0.60 | 0.87 | 2.0 | 78 | 0.43 | 0.52 | 0.23 | 0.19 | 0.78 |
| TP3-25-1310 | 25 | 2013-08-22 | 2013-10-27 | 11.7 | 11.4 | 12.0 | 0.36 | 0.46 | 0.79 | 2.1 | 102 | 0.49 | 0.48 | 0.20 | 0.18 | 0.79 |

| | | | | | | | | | | | | | | | | |
|---|---|---|---|---|---|---|---|---|---|---|---|---|---|---|---|---|
| TP3-25-1401 | 25 | 2013-10-27 | 2014-01-17 | 11.6 | 9.5 | 13.1 | 0.40 | 0.38 | 0.59 | 2.3 | 101 | 0.39 | 0.40 | 0.15 | 0.12 | 0.79 |
| TP3-25-1404 | 25 | 2014-01-17 | 2014-04-28 | 9.5 | 9.0 | 11.3 | 0.40 | 0.42 | 0.61 | 2.2 | 92 | 0.41 | 0.39 | 0.15 | 0.11 | 0.78 |
| TP2-35-1212 | 35 | 2012-10-25 | 2012-12-29 | 9.7 | 9.5 | 10.2 | 0.37 | 0.53 | 0.64 | 2.2 | 136 | 0.55 | 0.45 | 0.07 | 0.09 | 0.91 |
| TP2-35-1302 | 35 | 2012-12-29 | 2013-02-27 | 9.1 | 8.7 | 10.2 | 0.40 | 0.43 | 0.53 | 2.3 | 112 | 0.63 | 0.50 | 0.08 | 0.09 | 0.92 |
| TP2-35-1304 | 35 | 2013-02-27 | 2013-04-18 | 9.0 | 8.8 | 9.2 | 0.36 | 0.44 | 0.69 | 2.2 | 117 | 0.50 | 0.44 | 0.15 | 0.15 | 0.81 |
| TP2-35-1306 | 35 | 2013-04-18 | 2013-06-19 | 9.6 | 9.1 | 10.3 | 0.33 | 0.50 | 0.75 | 2.1 | 154 | 0.44 | 0.44 | 0.17 | 0.14 | 0.79 |
| TP2-35-1308 | 35 | 2013-06-19 | 2013-08-22 | 9.5 | 9.4 | 10.2 | 0.35 | 0.46 | 0.70 | 2.2 | 106 | 0.46 | 0.46 | 0.16 | 0.13 | 0.81 |
| TP2-35-1310 | 35 | 2013-08-22 | 2013-10-27 | 9.6 | 9.5 | 9.7 | 0.37 | 0.32 | 0.66 | 2.2 | 116 | 0.53 | 0.48 | 0.12 | 0.11 | 0.86 |
| TP2-35-1401 | 35 | 2013-10-27 | 2014-01-17 | 9.8 | 9.5 | 10.4 | 0.41 | 0.27 | 0.58 | 2.3 | 111 | 0.49 | 0.39 | 0.09 | 0.08 | 0.87 |
| TP2-35-1404 | 35 | 2014-01-17 | 2014-04-28 | 9.2 | 8.9 | 10.0 | 0.42 | 0.34 | 0.55 | 2.3 | 96 | 0.42 | 0.39 | 0.14 | 0.12 | 0.79 |
