## Supplementary material for "Quantifying spatiotemporal decoupling of GDGT-temperature relationships in a deep alpine lake": main text table

Table 1 GDGT-based proxies and equations used for calculation.

| Proxies | Equations | Reference |
|---|---|---|
| $TEX_{86}$ | $\frac{GDGT-2+GDGT-3+Cren'}{GDGT-1+GDGT-2+GDGT-3+Cren'}$ | Schouten et al. (2002) |
| BIT | $\frac{Ia+IIa+IIIa+IIa'+IIIa'}{Ia+IIa+IIIa+IIa'+IIIa'+Cren}$ | Hopmans et al. (2004) |
| GDGT-0/Cren | $\frac{GDGT-0}{crenarchaeol}$ | Blaga et al. (2009) |
| RI | $\frac{0\times GDGT-0+1\times GDGT-1+2\times GDGT-2+3\times GDGT-3+4\times Cren+4\times Cren'}{GDGT-0+GDGT-1+GDGT-2+GDGT-3+Cren+Cren'}$ | Zhang et al. (2016) |
| Cren/Cren' | $\frac{crenarchaeol}{crenarchaeol'}$ | Li et al. (2016) |
| CBT | $-\log(\frac{Ib+IIb+IIb'}{Ia+IIa+IIa'})$ | Weijers et al. (2007) |
| $MBT'_{5ME}$ | $\frac{Ia+Ib+Ic}{Ia+Ib+Ic+IIa+IIb+IIc+IIIa}$ | De Jonge et al. (2014a) |
| $MBT'_{6ME}$ | $\frac{Ia+Ib+Ic}{Ia+Ib+Ic+IIa'+IIb'+IIc'+IIIa'}$ | De Jonge et al. (2014a) |
| Index 1 | $\log(\frac{Ia+Ib+Ic+IIa'+IIIa'}{Ic+IIa+IIc+IIIa+IIIa'})$ | De Jonge et al. (2014a) |
| $IR_{6ME}$ | $\frac{IIa'+IIb'+IIc'+IIIa'+IIIb'+IIIc'}{IIa+IIb+IIc+IIa'+IIb'+IIc'+IIIa+IIIb+IIIc+IIIa'+IIIb'+IIIc'}$ | De Jonge et al. (2014b) |
